# TH2BS12P histone mark is enriched in the unsynapsed axes of the XY body and predominantly associates with H3K4me3-containing genomic regions in mammalian spermatocytes

**DOI:** 10.1101/455659

**Authors:** Iyer Aditya Mahadevan, Satyakrishna Pentakota, Raktim Roy, Utsa Bhaduri, Manchanahalli R. Satyanarayana Rao

## Abstract

Various studies have focussed on understanding the repertoire and biological function of the post-translational modifications that occur on testis-specific histone variants like TH2B, Transition Proteins etc. In our attempt to decipher the unique functions of histone variant TH2B, we discovered a new modification Serine 12 phosphorylation on TH2B (TH2BS12P) in spermatocytes. Our present study is aimed at understanding the function of the TH2BS12P modification in the context of processes that occur during meiotic prophase I. Immunofluorescence studies revealed that TH2BS12P histone mark is enriched in the unsynapsed axes of the sex body and is associated with XY body axes associated proteins like Scp3, γH2AX, pATM, ATR etc. We also observe that TH2BS12P is associated with DSB initiator Spo11 and with several recombination related proteins like pATM, ATR, Rad51, γH2AX etc *in vivo*. This modification was also found to associate with transcription and recombination related histone marks like H3K4me3 and H3K36me3 in the context of mononucleosomes. Genome-wide occupancy studies as determined by ChIP sequencing experiments revealed that TH2BS12P is localised to subset of recombination hotspots, but majorly associated with H3K4me3 containing genomic regions like gene promoters. Mass spectrometry analysis of proteins that bind to TH2BS12P containing mononucleosomes revealed many proteins linked with the functions of pericentric heterochromatin, transcription and recombination related pathways. We propose that TH2BS12P modification could act alone or in concert with other histone marks for recruitment of appropriate transcription or recombination protein machinery at specific genomic loci. This is the first report documenting the role of a post-translational modification of a germ cell specific histone variant in meiotic prophase I related events.

## Introduction

Mammalian spermatogenesis offers an excellent model system to study chromatin remodeling by histone variants as the testis is known to express a large number of core and linker histone variants in a stage-specific manner^28,54,55,61,63,^. It is believed that chromatin locus-specific histone replacement with histone variants could be a possible basis for genome reprogramming in germ cells. Testis-specific histone variants play a critical role during the germ cell development. H3t (testis-specific histone variant of H3) is essential for the process of spermatogonial differentiation and ensures entry into the meiosis^80^. The loss of TH2A and TH2B leads to male sterility with defects in cohesin release during interkinesis and histone to protamine replacement, suggesting an essential role of the testis-specific histone variants during critical periods of male germ development^68^.

TH2B was one of the earliest histone variants discovered in mammalian cells^27,38^. To date, it is the only testis-specific histone variant known to replace a core histone on a genome-wide scale replacing nearly 80-85% of H2B in spermatocytes and spermatids^41^. The pachytene nucleosome core particle harboring the TH2B protein was shown to be less compact compared to H2B containing nucleosome core particle^56,57^. hTSH2B (human TH2B) reconstituted histone octamer was found to be less stable than the H2B reconstituted histone octamer *in vitro*^32^. On the other hand, the loss of mouse TH2B is compensated for upregulation of H2B and compensatory histone modifications on the core histones H2B, H3 and H4 in germ cells. However, the expression of tagged-TH2B created a dominant negative phenotype, leading to defective histone to protamine replacement and ultimately, male sterility^41^. TH2B shares 85% sequence similarity with the canonical histone H2B. Since the major amino acid differences between TH2B and H2B lie in the amino terminal tail, we surmised that these differences or the post-translational modifications acquired by some of the residues could contribute to unique functions of TH2B. In this direction, Pentakota *et al*^51^ characterized the repertoire of post-translational modifications (PTMs) that occur on histone variant TH2B isolated from spermatocytes and spermatid stages. By computational analysis, it was also shown that the amino acid differences and the post-translational modifications acquired by some of the residues cause the destabilization of TH2B-containing nucleosomes. Histone PTMs are proving to be key molecular players in epigenomic functions^25,74^. Recently, various studies have focussed on understanding the post-translational modifications on testis-specific histone variants like TH2B^33,51^, TP1^18^, TP2^18^, HILS1^40^, etc.

During prophase I of meiosis, the homologous chromosomes synapse and undergo recombination at non-randomly selected loci. The exchange of genetic material is critical for the generation of diversity in the offspring. During leptotene interval, the global induction of Spo11 mediated DSBs occur throughout the genome triggering the DNA damage response (DDR)^7,42^. Subsequently, MRN (Mre11-Rad50-Nbs1) complex recruits ATM kinase, catalyzes the first level of H2AX phosphorylation to form γH2AX^6,11,29^. The end resection and strand invasion mediated by MRN complex, RAD51, DMC1 and other proteins are characteristic of the next stage, the zygotene interval. During the pachytene stage, BRCA1 senses asynapsis and recruits ATR kinase for amplification of DDR signals along the unsynapsed axes for the establishment of the γH2AX domain in the XY body^8,60,77^. The region of homology between the X and the Y chromosomes termed as pseudo-autosomal region (PAR), is limited in size (~800 kb in mice) and are largely unsynapsed during meiosis. Therefore, to ensure chromosome segregation with at least one crossover in PAR, a higher crossover density and increased DSB occurrence is observed in the PAR compared to that of autosomes^5^. The increase in DSB sites is caused by specialised chromatin configuration in PAR where DNA is organized on a longer axis with shorter chromatin loops compared to autosomes^23,30^. The non-PAR regions of the X and Y chromosomes are transcriptionally silenced during meiotic prophase by a process termed as meiotic sex chromosome inactivation (MSCI)^78,86^. The crossover products are generated during the subsequent diplotene stage. The completion of meiosis I yield secondary spermatocytes which undergo meiotic II division to produce haploid round spermatids.

In our efforts to gain insight into the unique functions of TH2B particularly during meiotic prophase I, we discovered a new post-translational modification in the amino terminal end; serine 12 phosphorylation on TH2B (TH2BS12P) which was previously not reported in the literature. Histone phosphorylation is linked to diverse biological processes like DNA damage and repair (DDR)^49^, apoptosis^46^ etc. In this study, we show that TH2BS12P modification is a histone mark associated with unsynapsed axes of the XY body in pachytene spermatocytes of rodents. This is also associated with chromatin domains interacting with proteins involved in transcription, meiotic recombination and DSB repair. ChIP-sequencing studies reveal that majority of TH2BS12P containing regions were not hotspot-related but associated with other H3K4me3 containing regions like gene promoters, enhancers etc. Additionally, this histone mark associated chromatin domains is associated with important proteins and histone marks linked to functions of gene regulation, meiotic recombination and pericentric heterochromatin formation in XY body, as revealed by mass spectrometry studies.

## Results

### TH2B Serine 12 phosphorylation (TH2BS12P) is a novel histone modification detected in mammalian spermatocytes-

Recently, we have described the various post-translational modifications of rat testicular TH2B isolated from spermatocyte and spermatids. By employing different set of procedures that includes enzyme digestion and post mass spectrometry analyses, we detected a new modification, TH2BS12P (TH2B serine 12 phosphorylation) which was not reported in our previous publication^51^. The representative MS/MS plot is shown in Fig 1A. The corresponding fragmentation table is in Fig S1A. As mentioned earlier, TH2B differs from H2B protein with majority of amino acid residues differing in the amino terminal tail. The structure of TH2B containing nucleosome model has been shown in Fig 1B. As expected, the serine 12 of TH2B is exposed to the solvent within the N-terminal tail. We also compared the N terminal amino acid sequences of TH2B across various mammalian species. As can be seen in Fig 1C, this serine residue is highly conserved. A comparison of N-terminal sequence of somatic H2B with TH2B, it can be seen that there is a serine residue at 14^th^ position, which is also conserved across all the somatic H2Bs among mammals (data not shown). The somatic H2BS14P modification has been shown to be involved in DNA repair processes in somatic cells^15^. Therefore, we were interested to delineate the function of the testis-specific H2B variant TH2B serine 12 phosphorylation in the context of DNA repair and meiotic recombination associated events in meiotic spermatocytes. For this purpose, polyclonal antibodies specific to this modification were generated in rabbits. The specificity of the antibody was determined by dot-blot, ELISA and western blotting analyses. Dot-blot assay validated the specificity of the antibody towards the TH2BS12P phosphopeptide but did not react with the H2B, H2BS14P and the unmodified TH2B peptides (Fig 1D). This was further substantiated by the ELISA assays wherein the antibody showed high reactivity towards the TH2B serine 12 phosphopeptide but not with the TH2B and H2B backbone peptides (Fig 1E). Using western blotting technique, the antibody showed reactivity towards testis nuclear lysates (Fig 1F, Lane ‘T’) but did not cross-react with the H2B containing liver nuclear lysate and recombinant TH2B protein (Fig 1F, lanes ‘L’ and ‘R’). Also, the reactivity of the antibody was abolished by preincubation with TH2B serine 12 phosphopeptide (Fig 1F, lane 2) but not with the TH2B unmodified peptide (Fig 1F, lane 3). The combination of mass spectrometry and western blotting analyses thus reveal that the TH2BS12P modification is physiological.

**Fig 1.**
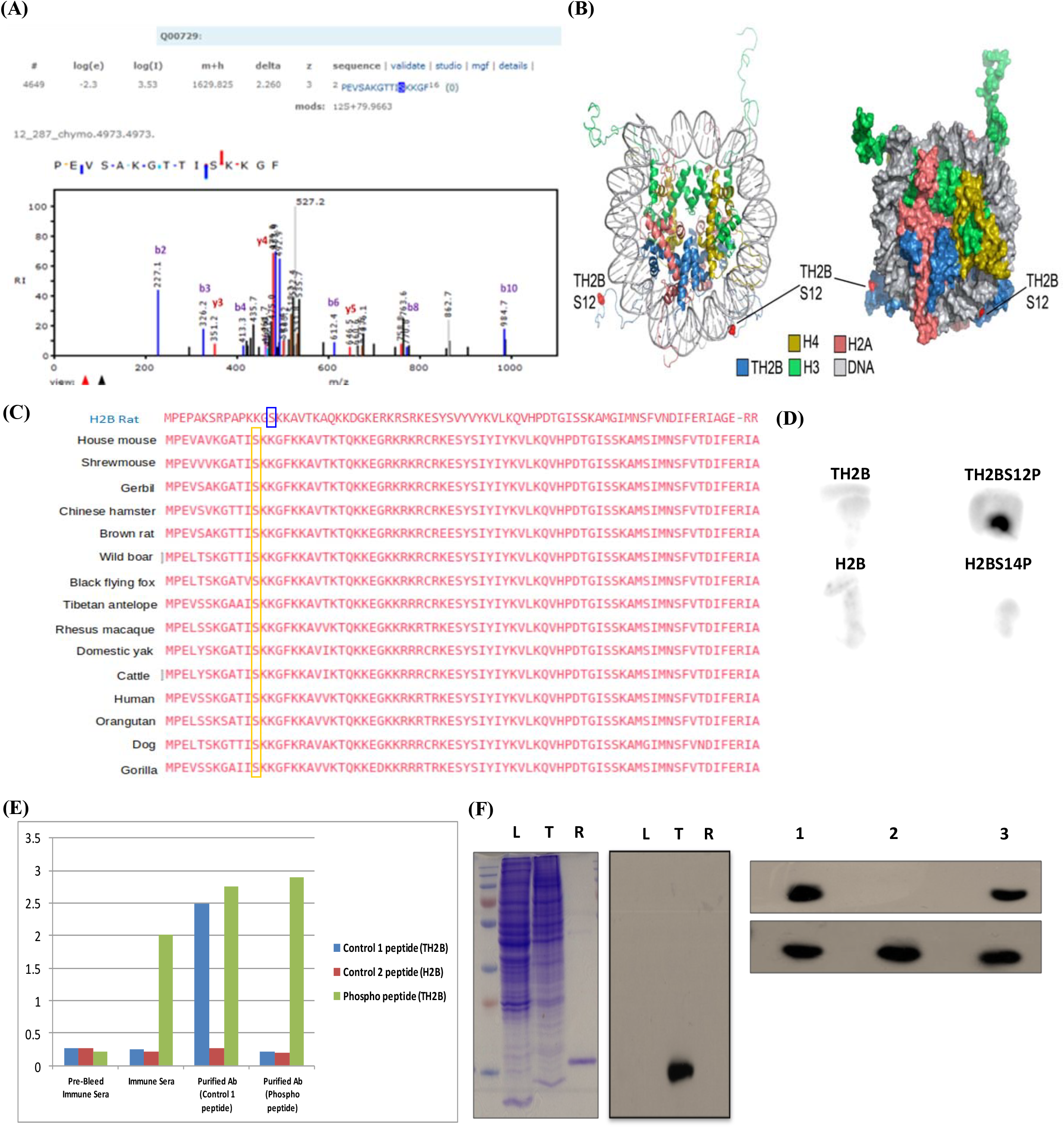
Identification of TH2BS12P modification by mass spectrometry and characterization of the TH2BS12P antibody. **(A)** Identification of TH2BS12P modification by LC-MS/MS technique. The y-axis indicates the relative intensity of MS/MS spectra and the x-axis indicates the mass to charge ratio. The phosphorylated serine residue of TH2B is highlighted in red. **(B)** Cartoon (left) and surface (right) representations of nucleosome consisting of TH2B, H2A, H3 and H4. Highlighted is the solvent-exposed residue (TH2BS12-red) that is phosphorylated. **(C)** Sequence comparison of TH2B histone variant in selected mammalian species- The alignment of TH2B proteins from various mammalian species was done using the Clustal Omega software, it shows the conservation of PTM site serine 12 residue in selected mammals. The selected species have been indicated on the left. H2B protein of rat has been given for reference at the top showing the neighboring serine 14 residue. The analogous residue of serine 12 of TH2B has been marked in blue box on H2B in the sequence alignment; **(D)** Dot-blot assay showing the specificity of the TH2BS12P antibody; peptides used are indicated near each spot. **(E)** ELISA-carried out with the Pre-bleed immune sera, Immune sera, TH2BS12P sera and the TH2BS12P purified antibody; blue bars represent reactivity of the mentioned sera or antibody with TH2B backbone peptide, red bars represent reactivity with H2B peptide and the green bars represent reactivity with the TH2B serine 12 phosphopeptide. **(F)** Western blotting analysis of endogenous TH2BS12P protein of rats in liver nuclear lysates, testis nuclear lysates and recombinant TH2B using the purified TH2BS12P antibody. L-Liver nuclear lysate, T-Testis nuclear lysate, R-recombinant TH2B. Coomassie stained gel is given for reference on the left. The panel on the right shows result of peptide competition assay-lane 1 represents no peptide control, lane 2 represents TH2BS12P antibody preincubated with TH2B serine 12 phosphopeptide (CKGTTI(pS)KKGFK), third lane TH2BS12P antibody preincubated with TH2B backbone peptide(CKGTTISKKGFK), western blot studies were carried out with the TH2BS12P antibody.

### TH2BS12P modification is densely localized to the axes of the XY body during the pachytene stage of meiosis prophase I-

After establishing the high specificity of the TH2BS12P antibody, we carried out immunofluorescence studies with the TH2BS12P antibody to examine the staining pattern during the meiotic prophase I. TH2B begins to express in preleptotene spermatocytes and continues to be present till the late stages of spermiogenesis^41^. Keeping this in mind, we carried out immunofluorescence staining of meiotic spreads of mouse testicular germ cells to examine whether the TH2BS12P modification is associated with any of the events that are characteristic of meiotic prophase I. We have used Scp3 as a marker to distinguish various stages of meiotic prophase I. We observe that TH2BS12P is detectable during the stages leptotene, zygotene and pachytene intervals as shown in Fig 2B. It is interesting to note that the backbone TH2B is detected all over the nucleus (Fig 2A), while TH2BS12P signal was more distributed as specific foci, suggesting a locus-dependent function. An important observation that is apparent in the staining pattern in pachytene spermatocytes is that TH2BS12P modification was found to be highly enriched at the axes of the XY body like structure (Fig 2B, pachytene spermatocytes). This was further corroborated by colocalization analysis of TH2BS12P signal with Scp3 as shown in Fig 2D (Scp3) and was highly enriched in the XY body in comparison to the leptotene, zygotene or the rest of pachytene nuclei. This was also confirmed in meiotic spreads of rat testicular cells where we observed increased enrichment of TH2BS12P signal in the XY body like structure (Fig S2A, pachytene). We therefore conjectured that this modification may have a XY body specific function in spermatocytes. At the same time, we observed many foci outside the sex body in pachytene cells suggesting the role of TH2BS12P may not be just restricted to XY body specific functions.

**Fig 2.**
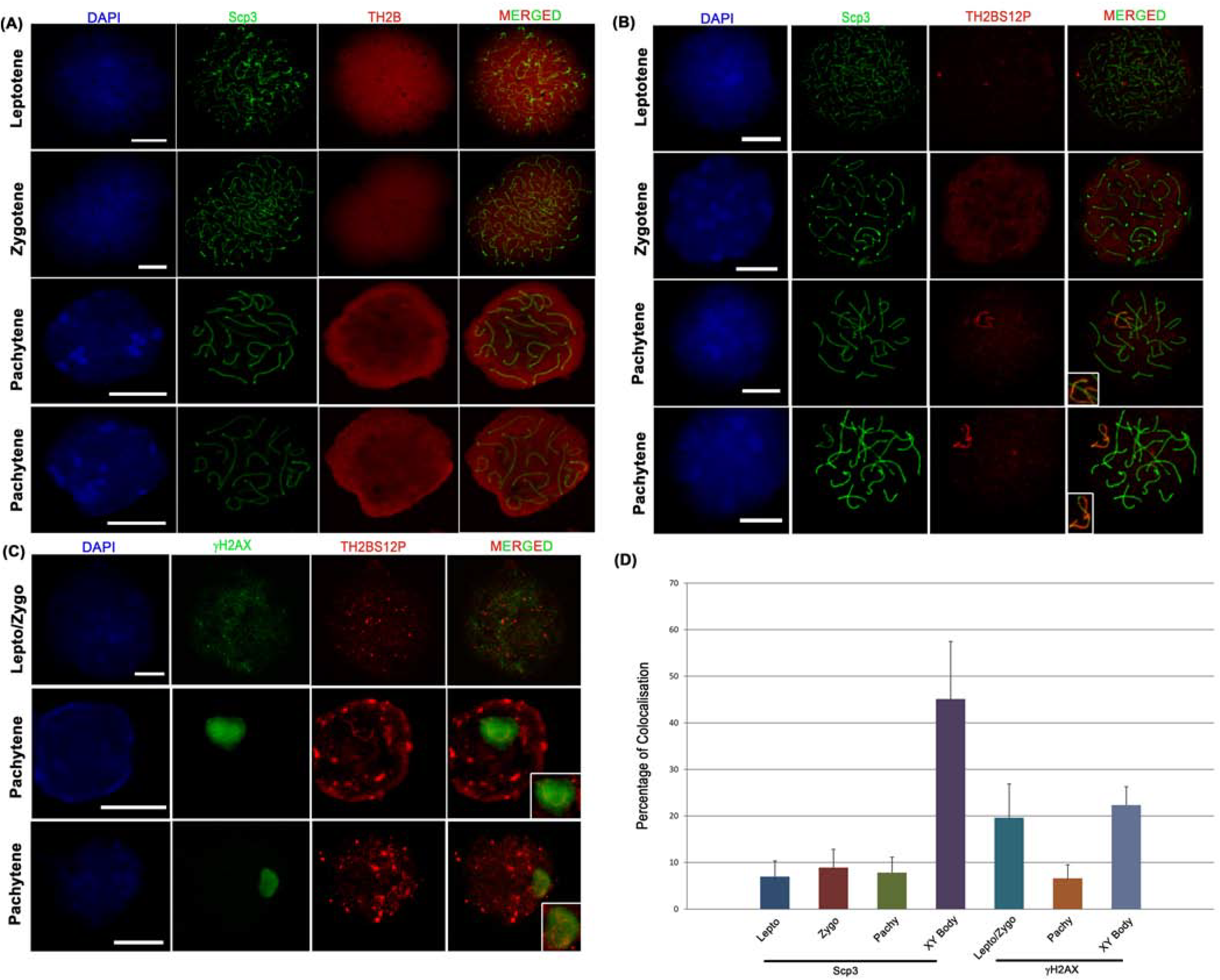
TH2BS12P modification is enriched in the axes of the XY body and highly colocalizes with γH2AX in leptotene/zygotene nuclei-. **(A)** Immunofluorescence studies of backbone TH2B and Scp3 in leptotene (1st panel), zygotene (2nd panel) and pachytene stages (3rd and 4th panels) of meiotic prophase I of mouse germ cells. Nuclei are stained with DAPI. The inset squares in panel 3 and 4 represents the XY body. **(B)** Colocalization studies of TH2BS12P modification with synaptonemal complex protein Scp3 across leptotene (1st panel), zygotene (2nd panel) and pachytene stages (3rd and 4th panels) of meiotic prophase I. The inset squares in panel 3 and 4 represents the XY body. **(C)** Colocalization studies of TH2BS12P with γH2AX in leptotene/zygotene (1st panel) and pachytene spermatocytes (2nd and 3rd panels). The inset squares in panel 3 and 4 represents the XY body. **(D)** Quantitation of colocalization percentages of TH2BS12P with Scp3 and γH2AX in leptotene (lepto), zygotene (zygo), pachytene and XY body (XY body). All data were confirmed with germ cells from at least three independent mice. The data are plotted as mean ± S.D. Scale bars, 10 μm.

H2AX is required for chromatin remodeling and sex chromosome inactivation in male meiosis^14^. We confirmed the enrichment and localization of this TH2BS12P modification in the XY body of the pachytene nucleus using the sex body specific marker γH2AX. As can be seen in Fig 2C, TH2BS12P colocalizes with γH2AX corresponding to the axes of the XY body in the pachytene spermatocytes. The degree of colocalization of TH2BS12P with γH2AX in the XY body was found to be high in the XY body (~20%) as compared to the whole pachytene spermatocyte nucleus (~5%) as indicated in Fig 2D. Apart from the role of γH2AX in chromatin remodeling in the XY body, ATM dependent phosphorylation of H2AX to form γH2AX, is the hallmark of autosomal DSB formation. γH2AX is associated with meiotic DSBs in both autosomes and sex chromosomes. On these lines, we were also interested to carry out colocalization experiments of TH2BS12P with γH2AX in the leptotene/zygotene intervals. We also observe a good degree of colocalization of TH2BS12P with γH2AX in the leptotene-zygotene nuclei (Fig 2A). We observe colocalization percentages of TH2BS12P with γH2AX in leptotene/zygotene nuclei (~20%) similar to that of XY body (~21%) (Figure 2D, γH2AX). The fact that TH2BS12P is enriched in the axes of the XY body and colocalizes with γH2AX in XY body and leptotene/zygotene intervals equally indicates that this histone mark may be associated with recombination and repair related proteins in spermatocytes.

In a previous study, H2BS14P was shown to stain the XY body of pachytene spermatocytes in mouse^15^. Since, our data also suggests that TH2BS12P to localize in the XY axes, we wondered whether the TH2BS12P antibody did crossreact with H2B or its posttranslational modifications in spermatocytes. However, on re-examination of the published data, we found that the same H2BS14P commercial antibody cross-reacts also with *in vivo* TH2B (Fig S1B, H2BS14P). The antibody has been withdrawn and no more available. Therefore, we generated a H2B specific antibody, validated its reactivity by dot-blot assay and western blotting with liver and testicular histones as shown in Fig S1C, H2B (Lanes ‘L’ and ‘T’). We carried out staging of the H2B antibody with Scp3 to obtain the staining pattern at various stages of meiotic prophase I. The staining of backbone H2B was found to be not intense contrary to that previously reported for H2BS14P modification in all the stages of meiotic prophase I (Fig S1D). Furthermore, H2B staining did not coincide with the XY body of the pachytene spermatocyte (Fig S1D, pachytene). Nevertheless, the combination of Western blotting with peptide competition, ELISA and dot-blot assays prove that the TH2BS12P antibody did not cross-react with any of the post-translational modifications on H2B (including H2BS14P modification). This negates the finding of the previous report where H2BS14P was shown to localize to the XY body suggesting that the reported staining pattern is possibly an artefact.

### TH2BS12P modification is densely localised in the unsynapased axes and associated with various recombination-related proteins like Spo11, pATM, ATR in the XY body of the pachytene spermatocyte-

During leptotene interval, initiation of meiotic recombination is followed by DSB formation, triggered by local phosphorylation of H2AX by pATM to form γH2AX^3,11,29^. From mid-zygotene interval, unsynapsed chromosomes are marked by ATR, where the latter carries out the second level of H2AX phosphorylation^8,60^. ATM-dependent signaling pathway of γH2AX formation is the hallmark of autosomal recombination during leptotene and zygotene, whereas the ATR-dependent signaling operates in the XY body specific formation of γH2AX foci during the pachytene stage^11,15,29,44^. ATR is a multifunctional protein required for proper loading of strand invasion proteins, elongation of synaptonemal complex and recruitment of recombination proteins players on unsynapsed axes^45,84^. Therefore, we next sought to perform colocalization studies of pATM and ATR proteins with TH2BS12P to examine whether TH2BS12P is involved in the ATM or ATR signaling pathways or both. To test this, we performed colocalization studies with pATM and ATR kinases again in leptotene, zygotene and pachytene intervals of meiotic prophase I. Figure 3A shows the colocalization of TH2BS12P with pATM across the different stages. We observe clear colocalization of TH2BS12P with pATM in the leptotene/zygotene stages. Upon further quantitative analysis, we observed high colocalization percentage of 62% in the leptotene/zygotene nuclei (Fig 3D, pATM). This suggests that this histone modification has an important role to play in autosomal DSB formation characteristic of leptotene/zygotene interval. Furthermore, we observe a high colocalization percentage of about 25% with pATM corresponding to the axes of the XY body as indicated in Fig 3B and 3D (pachytene). In the case of the pachytene spermatocyte, we observed a high colocalization of TH2BS12P with pATM in the XY body compared to rest of the pachytene nucleus (Fig 4D, pachytene & XY body).

**Fig 3.**
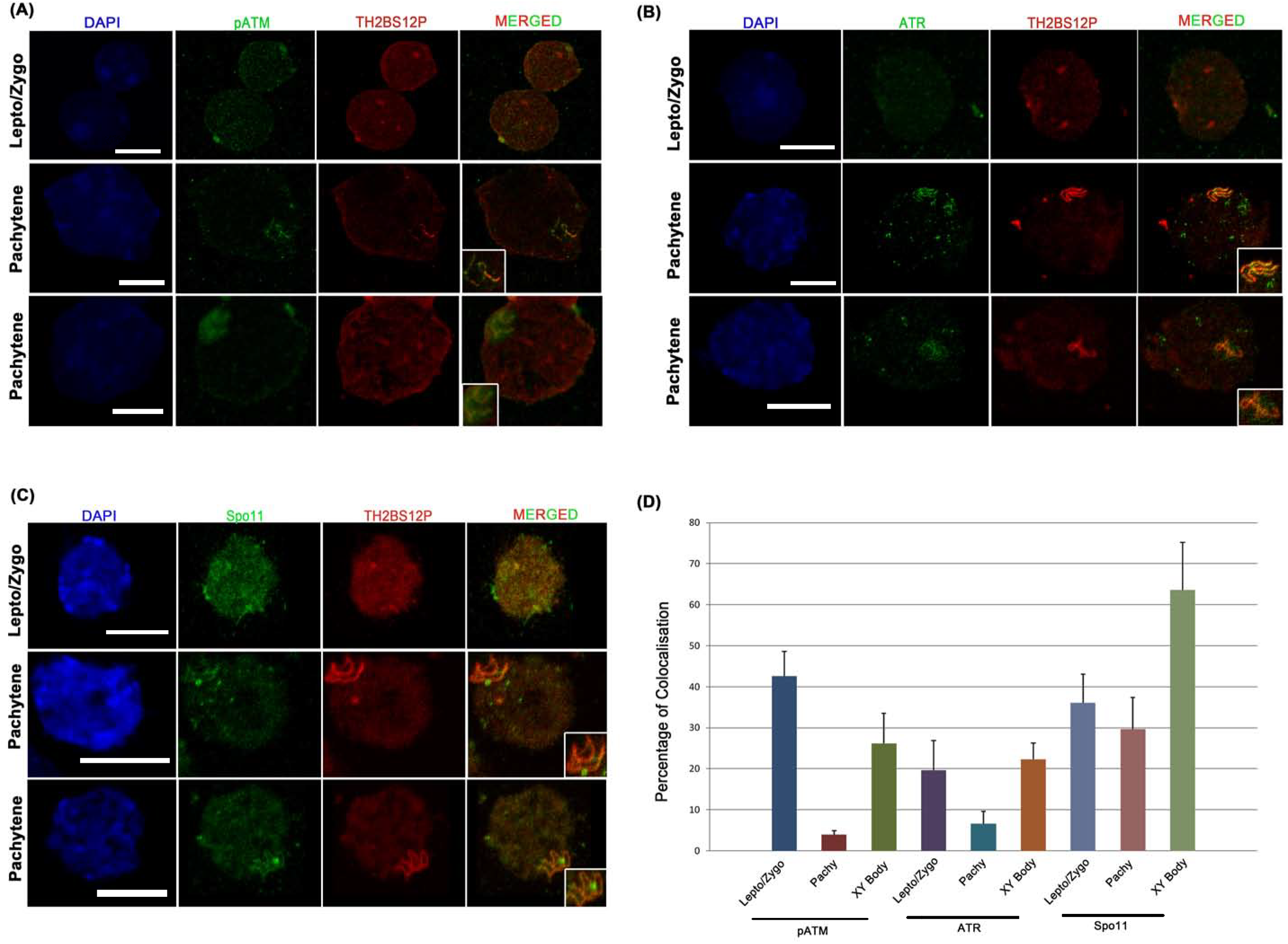
TH2BS12P colocalizes with DSB initiator Spo11 and is highly enriched in the unsynapsed axes of the XY body of the pachytene spermatocyte. **(A)** Colocalization studies of TH2BS12P with pATM in leptotene/zygotene (1^st^ panel) and pachytene spermatocytes (2^nd^ and 3^rd^ panels); **(B)** Colocalization studies of TH2BS12P with ATR kinase in leptotene/zygotene (1^st^ panel) and pachytene spermatocytes (2^nd^ and 3^rd^ panels) **(C)** Colocalization studies of TH2BS12P and Spo11 in leptotene/zygotene (1^st^ panel) and pachytene spermatocytes (2^nd^ and 3^rd^ panels). The inset squares in 2^nd^ and 3^rd^ panels of all the figures from A-C shows the XY body. Nuclei were visualized by DAPI staining. Scale bars, 10 μm. **(D)** Quantitation of colocalization percentages of TH2BS12P with Spo11, pATM and ATR across the selected intervals of meiotic prophase I: leptotene (lepto), zygotene (zygo) and pachytene with XY body (XY body). All data were confirmed with germ cells from at least three independent mice. The data are plotted as mean ± S.D.

**Fig4.**
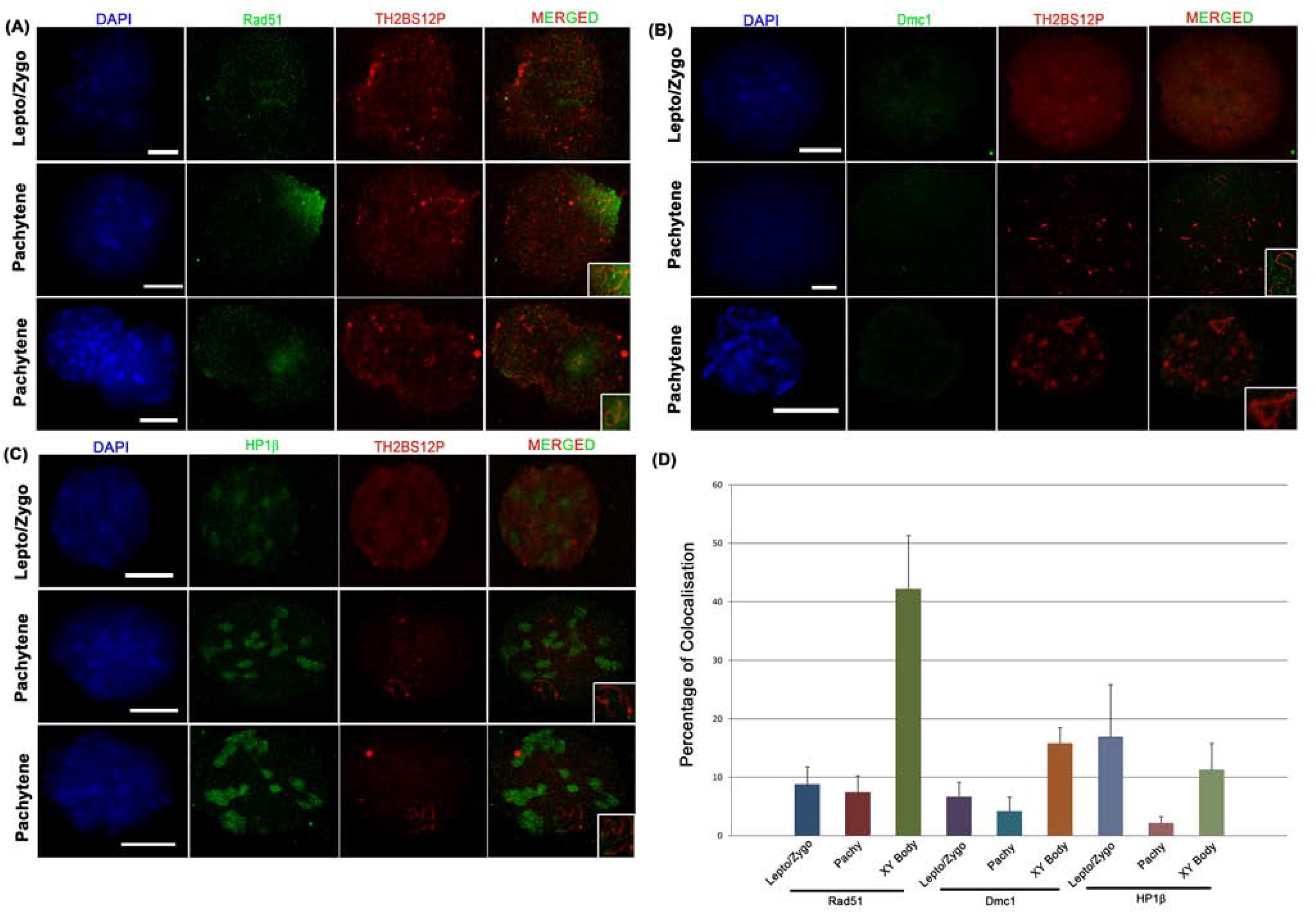
TH2BS12P is associated with recombinases Rad51 and Dmc1 and heterochromatin related protein HP1β in spermatocytes-. **(A)** Colocalization studies of TH2BS12P with Rad51 in leptotene/zygotene (1st panel and pachytene stages (2^nd^ and 3^rd^ panels) of meiotic prophase I of mouse germ cells. **(B)** Colocalization studies of TH2BS12P and Dmc1.**(C)** Immunofluorescence studies of TH2BS12P and HP1β in leptotene/zygotene (1st panel) and pachytene spermatocytes (2^nd^ and 3^rd^ panels) of mouse testicular cells. The inset squares in panel 2 and 3 of all the figures represents the XY body. Nuclei were visualized by DAPI staining. Scale bars, 10 μm. **(D)** Quantitation of colocalization percentages of TH2BS12P with Rad51, Dmc1 and HP1β across various stages of meiotic prophase I: leptotene (lepto), zygotene (zygo) and pachytene with sex body (XY body). All data were confirmed with germ cells from at least three independent mice. The data are plotted as mean ± S.D.

Colocalization studies were also carried out of TH2BS12P and ATR to determine whether TH2BS12P is involved in the signaling events related to XY body recombination. While we observe a good colocalization of TH2BS12P with ATR protein in the leptotene/zygotene nuclei, there was ~22% distinct colocalization seen specifically in the axes of the XY body as seen in Fig 3B. We observed a higher degree of colocalization with ATR in the XY body compared to leptotene/zygotene nuclei in agreement with the above data (Fig 3D, ATR). On the basis of colocalization of TH2BS12P with ATR and pATM in the XY axes during the pachytene interval, we conclude that this modification is indeed localised to the unsynapsed axes of the XY body. Altogether, these data show that TH2BS12P is highly associated with pATM kinase in leptotene/zygotene nuclei and associated with both pATM and ATR kinases in the XY body. The above results establish a role of TH2BS12P in the dynamics of autosomal and XY body recombination during the meiotic prophase I.

XY body harbors various proteins related to recombination and heterochromatin formation^19^. DDR proteins that accumulate in the XY body, have different and distinct localisation patterns. Proteins like γH2AX are spread over the entire sex chromosomes (axes + loops), whereas ATR, pATM, Rad51 are localised in the unsynapsed axes. This suggests that unsynapsed axes are different from other chromosomal regions of the sex body in terms of distinct localisation patterns of various DDR related proteins^35^. Since pATM and ATR are marker proteins of the unsynapsed axes and colocalize with TH2BS12P, we conclude that TH2BS12P histone mark is enriched in the unsynapsed axes of the XY body of pachytene spermatocyte.

Spo11 dependent DNA breaks are involved in the initiation of meiotic recombination^10,24,37^. Spo11 is also essential for sex body formation, the disruption of which leads to male infertility^6^. Keeping this in mind, we carried out colocalization studies of TH2BS12P modification with Spo11 and also quantified colocalization percentages in all the intervals of meiotic prophase I. TH2BS12P colocalizes with Spo11 in the leptotene/zygotene nuclei as shown in Fig 3C. In agreement with this finding, we observed a high colocalization percentage of TH2BS12P with Spo11 in the leptotene-zygotene nuclei (~34)[Fig 3D, Spo11]. In addition, Fig 3C (pachytene spermatocytes) clearly demonstrates the colocalization of TH2BS12P with Spo11 in the axes of the XY body. This observation is consistent with the colocalization analysis which demonstrates that TH2BS12P highly colocalizes with Spo11 in the axes of the XY body (~62% in average) compared to the whole pachytene spermatocyte (~30 % in average) [Fig 3D, Spo11]. Altogether, we observed significant colocalization of Spo11 with TH2BS12P in leptotene/zygotene nuclei and in the axes of XY body of pachytene spermatocyte. These results therefore confirm that TH2BS12P modification is associated with Spo11 that is involved in DSB initiation in meiotic stages of autosomal and XY body recombination.

Homologous recombination is mediated by two important proteins Rad51 and meiosis-specific Dmc1. They share high degree of amino acid sequence identity and exhibit similar catalytic activities *in* vitro^69,76^. We were interested to carry out colocalization experiments of TH2BS12P with Rad51 and Dmc1 to determine whether this histone mark is associated with strand invasion processes characteristic of the zygotene interval.

TH2BS12P showed minimal colocalization with Rad51 in the leptotene/zygotene and pachytene spermatocyte as depicted in Fig 4A. This was supported by colocalization analysis which revealed low degree of colocalization of TH2BS12P with Rad51 in the leptotene-zygotene or pachytene nuclei (Fig 4D, Rad51). Instead, TH2BS12P was found to highly colocalize Rad51 in the unsynapsed axes of the XY body during the pachytene interval (Fig 4A, pachytene). Quantitatively, colocalization percentages for TH2BS12P and Rad51 were about ~7% and ~41% average in pachytene spermatocytes and XY body respectively. Altogether, these data show that TH2BS12P is associated with Rad51 in the axes of the XY body (Fig 4D, Rad51). Similarly,

TH2BS12P was not found to colocalize with Dmc1 to a significant extent in leptotene, zygotene, and pachytene intervals (Fig 4B; Fig 4D, Dmc1). Also, we observed highest colocalization percentage of TH2BS12P with Dmc1 in the XY body (~15%) compared to that in leptotene, zygotene and pachytene spermatocytes (Fig 4D, Dmc1). These led to the conclusion that TH2BS12P containing nucleosomes were also associated with repair related proteins in the XY body, not to that extent in the case of autosomal DSBs. Homologous chromosomes are completely synapsed at pachytene interval. Since heterochromatin proteins like HP1β are localized in the XY body during pachytene interval, we were curious to examine the colocalization of TH2BS12P with HP1β to determine whether TH2BS12P histone mark has any role in heterochromatin formation in the sex body. HP1β and HP1γ decorate the whole sex body during the pachytene interval and are involved in stabilisation of heterochromatin state in the XY body^34,75^. In addition, they are also necessary for proper chromosome segregation. We did observe that the colocalization of TH2BS12P with HP1β was much less in all stages of meiotic prophase I (Fig 4C). Quantitation analysis also revealed only significant colocalization with HP1β in the leptotene/zygotene and XY body than in pachytene spermatocytes (Fig 4D, HP1β). This led us to conclude that TH2BS12P was primarily associated with heterochromatin formation during the meiotic prophase I. Taken together, the immunofluorescence studies reveal that the TH2BS12P modification is highly associated with the proteins related to DNA recombination and repair processes in spermatocytes. We have also repeated the colocalization experiments of TH2BS12P in meiotic spreads prepared from 25-30 day old rat testes. These immunofluorescence images are given in Supplementary figures S2 and S3 which reveal that similar pattern is observed in rat meiotic spreads.

### TH2BS12P is associated with proteins associated with DSB initiation and meiotic recombination

Meiotic DSBs do not occur randomly in the genome, but occur at specific sites marked by histone marks H3K4me3 and H3K36me3, which provides a suitable chromatin template for the Spo11 mediated DSB formation^53,71^. To confirm the immunofluorescence studies, we were interested in carrying out studies to determine the association of TH2BS12P with important marker proteins related to DSB initiation protein like Spo11 and Strand Invasion protein like Rad51 using co-immunoprecipitation assays. To shed light on whether TH2BS12P is associated with Spo11 and the associated histone marks, we carried out ChIP analysis where Spo11 associated chromatin complexes were immunoprecipitated using a Spo11 antibody and scored for TH2BS12P and H3K4me3 by western blotting. From Fig 5A ((i) Spo11), it can be seen that Spo11 interacts with TH2BS12P containing chromatin domains in spermatocytes. Also, Spo11 co-immunoprecipitates with H3K4me3 histone mark consistent with the literature where H3K4me3 is associated with initiation of DSB formation (Fig 5A (i)). This finding strongly supports the association of Spo11 with TH2BS12P containing nucleosomes.

**Fig 5.**
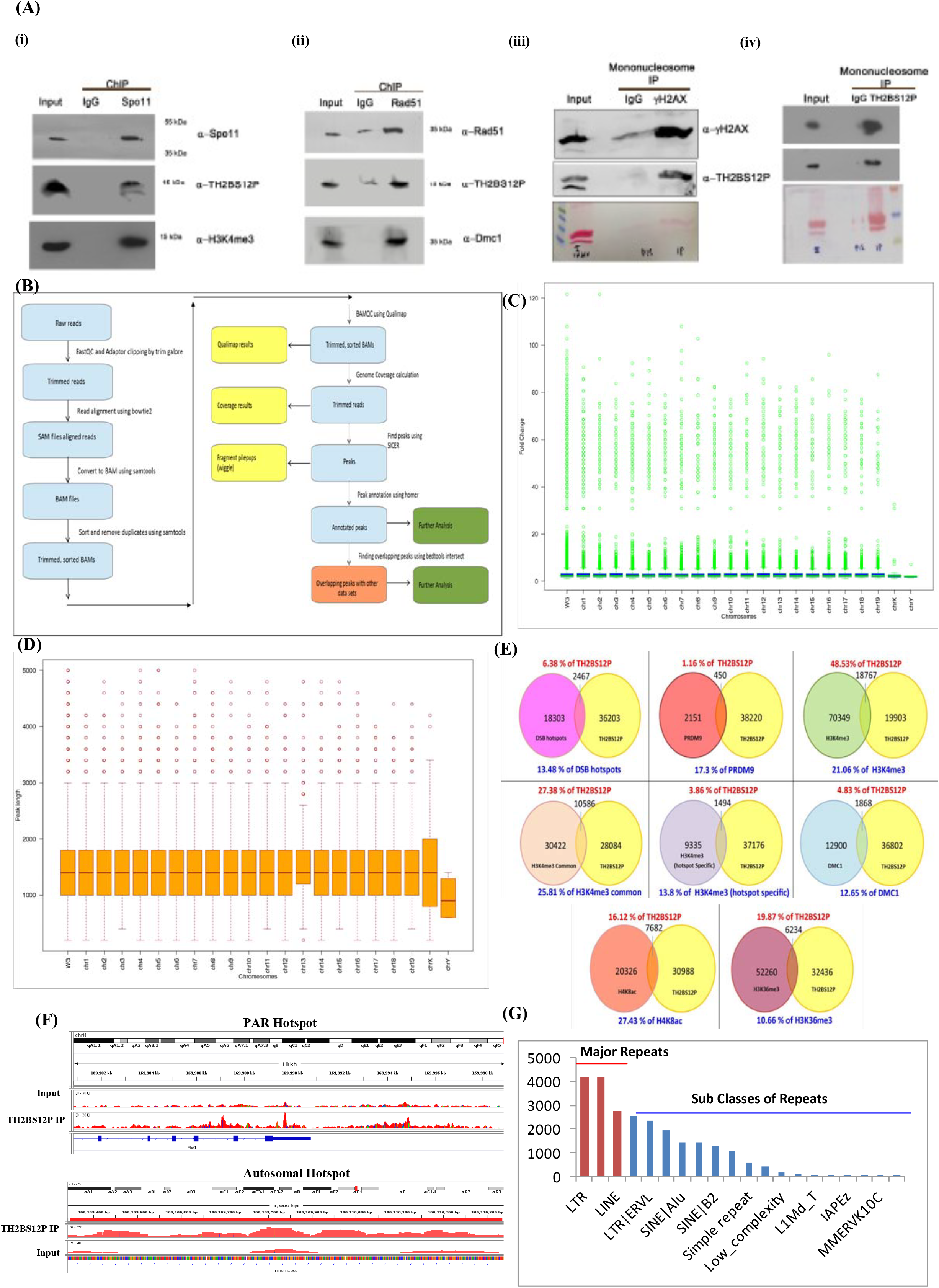
Co-immunoprecipitation of TH2BS12P with recombination associated proteins and Genome-wide occupancy of TH2BS12P in P12 mouse testicular cells. **(A)** Spo11and Rad51 are associated with chromatin fragments containing TH2BS12P histone mark-Chromatin Immunoprecipitation studies with **(i)** anti-Spo11 and **(ii)** anti-Rad51 using testes of 20-25 day old rats, followed by detection of co-immunoprecipitation by western blotting using the respective antibodies shown in alpha (α) alongside the blot. **(iii)** and **(iv)** TH2BS12P containing mononucleosomes coexist with DSB initiator mark γH2AX-Mononucleosomes IP studies determining the coexistence of histone marks with TH2BS12P with γH2AX; alpha alongside the blot indicates the antibodies used for blotting. Ponceau stained blots have been given for reference. The numbers represent molecular weight in kilodaltons. The first lane in all the blots represent input fraction, 2nd lane IgG elute fraction and 3rd lane immunoprecipitation reactions with the mentioned antibodies; **(B)** Workflow of ChIP seq analysis for TH2BS12P; **(C)** Chromosome-wise distribution of fold enrichment of the TH2BS12P peaks; **(D)** Chromosome-wise distribution of peak-length of TH2BS12P peaks; **(E)** Pie chart displaying the percentage overlap of TH2BS12P histone mark with the ChIP seq datasets available for various proteins and histone marks related to meiotic recombination and transcriptional regulations like DSB hotspots, PRDM9, H3K4me3, H3K4me3 common (TSS, enhancers), B6 specific H3K4me3 (hotspots), Dmc1, H4K8ac and H3K36me3 peaks; **(F)** TH2BS12P in mappable PAR and autosomal DSB hotspot regions-Figure of IGV Genome browser showing the enrichment of TH2BS12P (IP track) compared to Input track in the PAR region of the X chromosome and in the autosomal hotspot region located in chromosome 5, the peaks in the input and IP track (both marked in red color) are shown in the same scale for comparison; **(G)** Localization of TH2BS12P histone mark at various repeat elements of the mouse genome-The major classes of repeats include LTR, LINE and SINE elements. The subclasses of these major repeats have also been represented.

Rad51 is expressed in both mitotic and meiotic cells, whereas the expression of Dmc1 is meiosis-specific, which is suggestive of their context specific roles *in vivo*. Rad51 and Dmc1 form the multiprotein recombination complex in interaction with Scp3 to mediate recombinational repair in spermatocytes^76^. An important question that arises then, is whether TH2BS12P containing nucleosomes are binding sites for strand invasion related protein Rad51. As can be seen from Fig 5A ((ii) Rad51), ChIP-western blot analysis revealed that TH2BS12P was associated with chromatin fragments containing Rad51 which is in agreement with our colocalization studies with Rad51. Dmc1 is known to interact with Rad51 in spermatocytes, therefore was used as a positive control for the ChIP reaction. The combination of immunofluorescence and chromatin immunoprecipitation studies supported that TH2BS12P histone mark is associated with major proteins belonging to the process of DDR and recombination in XY body of mammalian spermatocytes.

As mentioned earlier, γH2AX is a marker for DSBs which appears on meiotic chromosomes in a Spo11-dependent fashion. We employed MNase digestion followed by immunoprecipitation to address whether TH2BS12P is physically associated with γH2AX in context of mononucleosomes. By IP assays, we prove that γH2AX pulled down TH2BS12P protein (Fig 5A, (iii) γH2AX). Conversely, γH2AX is also found in TH2BS12P elute fraction (Fig 5A, (iv) TH2BS12P). Together, we prove by immunoprecipitation assays that TH2BS12P is associated with Spo11, Rad51 and γH2AX. Therefore, biochemical assays confirm the results obtained from immunofluorescence data proving the association of TH2BS12P in DSB repair and meiotic recombination associated phenomena.

### Genome-wide occupancy of TH2BS12P-

Since meiotic recombination is a non-random event and occurs at specific loci termed as meiotic recombination hotspots, we were curious to examine the possible association of TH2BS12P at these hotspot loci. Since the genetics of homologous recombination and hotspot sequences are well characterised in C57BL6 mouse species, we performed ChIP-seq analyses of genome wide distribution of TH2BS12P to obtain a comprehensive picture of the association of TH2BS12P with genomic regions in leptotene/zygotene cells to understand the proportion of this modification in context of autosomal DSBs. ChIP seq of TH2BS12P was carried out in P12 testicular cells, since the mouse recombination hotspots as well as the ChIP seq data of many of recombination and transcription related proteins have been generated using P12 mouse testicular cells. The workflow of the ChIP sequencing protocol and computational analyses is given in Fig 5B. After measuring the chromosome-wise fold change distribution, it was observed that fold change tends to be uniform across all the chromosomes (Fig 5C). A similar pattern has also been observed while taking the quantitative measurement of chromosome-wise peak length distribution of peaks identified by SICER as mentioned in methods section (Fig 5D).

### TH2BS12P is localised to subset of recombination hotspots and highly associated with H3K4me3 containing genomic regions like gene promoters in mouse spermatocytes

Recombination hotspots are defined as genomic regions with high enrichment of PRDM9 catalysed H3K4me3 and H3K36me3 marks^4,21,47,48,53,71^. We carried out the analyses using Bedtools Intersect tool to decipher the extent of association of TH2BS12P with the reported autosomal and XY body related hotspots. Surprisingly, only about 13.48% of published DSB hotspots were found to overlap with TH2BS12P binding sites (Fig 5E, DSB Hotspots). On the other hand, ChIP seq studies revealed that only 6.38% of TH2BS12P peaks were found to be associated with the known recombination hotspots (Fig 5E, DSB Hotspots). The analysis also revealed that about 17.3% of PRDM9 peaks were found to correlate with TH2BS12P containing genomic loci (Fig 5E, PRDM9). Taken together, these results show that TH2BS12P is localised to subset of total recombination hotspots in mice. Further, we carried out similar analyses to further understand the overlap of TH2BS12P with other recombination and transcription related proteins and histone marks.

We observed an interesting phenomenon, where about 48.5% of TH2BS12P peaks were found to coincide with H3K4me3 modification (Fig 5E, H3K4me3).

Recombination related H3K4me3 marks are distinct from the transcription related H3K4me3 marks^71^. Apart from their presence at recombination hotspots, H3K4me3 marks are also present in open chromatin regions like TSS, enhancers etc^62,73^. Next, we were interested in determining the extent of association of TH2BS12P histone in transcription or recombination related processes, by overlapping with specific H3K4me3 datasets. ‘H3K4me3 common’ peaks indicate the peaks common to C57BL6 and RJ2 species, indicates the regulatory elements of the genome such gene promoters, enhancers etc^17^. We found 25.81% of ‘common H3K4me3’ peaks associated with TH2BS12P mark. Conversely, 27.38% of TH2BS12P peaks were associated with ‘H3K4me3 common’ marks (Fig 5E, H3K4me3 common). This gives a clear indication that majority of association of TH2BS12P histone mark is towards other open chromatin regions like gene promoters, enhancers etc. ‘B6 specific’ H3K4me3 marks refer to the hotspot specific H3K4me3 marks in C57BL6 mouse species^17^. A minor percentage of 13.8% of ‘B6 specific’ H3K4me3 marks and 12.65% of total Dmc1 peaks is associated with TH2BS12P, reinforcing the conclusion that the majority of localisation of TH2BS12P occurs towards H3K4me3 regions that exclude the hotspots (Fig 5E, H3K4me3 hotspot specific data). Furthermore, H4K5 and K8 acetylation marks are hallmarks of active gene promoters in mammalian spermatocytes^16^. To better understand the relationship of TH2BS12P histone mark with active gene promoters, we carried out similar analyses with H4K8ac mark using Bedtools Intersect. We found 27.43% of total H4K8ac mark containing genomic regions were associated with TH2BS12P histone mark, suggesting a gene regulatory function of TH2BS12P in spermatocytes (Fig 5E, H4K8ac).

In addition to catalysing H3K4 trimethylation, PRDM9 also catalyses H3K36me3 formation. But in case of TSS, SETD2 is the major histone methyltransferase that could catalyse H3K36 trimethylation^53^. H3K36me3 is predominantly localised in the gene bodies in testicular cells. The co-association between H3K36me3 and TH2BS12P was restricted to around 19.87% of total TH2BS12P bound loci. On the other hand, 10.66% of total H3K36me3 mark is related to TH2BS12P mark (Fig 5E, H3K36me3). This could mean that the association of TH2BS12P with H3K36me3 occurs at recombination hotspots and gene bodies. Although, we found a good degree of colocalization with important recombination related proteins both in the autosomal and XY body regions, the proportion of occupancy of TH2BS12P were found to be about 13.48% of total hotspot regions. To conclude, ChIP seq analyses further reveal that majority of TH2BS12P loci were not hotspot-related but are closely related to H3K4me3 associated regions like gene promoters, enhancers etc.

Recombination in the XY body is restricted to a small region of homology between the X and Y chromosome, that is called the PAR region. This region is highly enriched with repetitive elements, so much that some of the published ChIP seq studies have not been able to map the entire region. However, the previous report had mapped a small non-repetitive region of PAR wherein all the recombination related marks like H3K4me3, H3K36me3 and Dmc1 were found to be present in the region^53^. Therefore, we analysed whether TH2BS12P is possibly associated with the reported mappable-PAR region. Confirming the XY body localisation of this modification and colocalization with other XY body proteins, ChIP seq in leptotene-zygotene cells revealed the association of this modification in the mappable PAR region. We found the TH2BS12P IP signal considerably higher than that of Input track, suggesting good enrichment of this modification in the PAR region, thus corroborating our findings of staining in the XY body as earlier shown by IF studies (Fig 5F, PAR hotspot). For representing TH2BS12P localisation at autosomal hotspots, we have shown the enrichment of TH2BS12P in IP track versus Input at the selected autosomal hotspot location in chromosome 5 (Fig F, autosomal hotspot).

It is important to note that the number of TH2BS12P peak-overlaps at recombination hotspots may be underrepresented for the reason that the hotspots are also associated with repetitive elements like L1(LINE1) elements, ERVK family LTR elements etc^87^. For determining the localisation of TH2BS12P histone mark at repeat elements, we further characterised the genomic regions of TH2BS12P and plotted the peak number with respect to number of annotated repeat elements in the mouse genome taken from UCSC table browser. From the surface overview, it has been observed that TH2BS12P peaks associate with the major classes of LTR, SINE and LINE elements (Fig 5G, major repeats). With close analyses, it has been identified among the subclasses of repeat elements, the localisation of TH2BS12P occurs at DSB-associated-elements like LINEL1, LTR ERVK suggesting that hotspots associated with TH2BS12P could indeed be underrepresented because of its localisation at the aforementioned repeat elements (Fig 5G, sub classes of repeats).

### TH2BS12P is predominantly localised at H3K4me3 containing genomic regions like gene promoters and interacts with TSS associated histone marks *in vivo*

To understand the interaction of TH2BS12P with important genomic regions in spermatocytes further, we used aggregation plots to determine the distance of TH2BS12P peaks with respect to the centre of binding sites of various recombination markers like PRDM9, H3K4me3 and transcription associated protein markers like H3K4me3, H3K36me3, etc. We used Aggregation and Correlation Toolbox to address this question, wherein ChIP seq signal intensities of TH2BS12P histone mark are averaged at user-defined anchor points (of protein and histone marks)^82^. As seen in peak overlap data, we found increased peak correlation of TH2BS12P with PRDM9 with respect to distance calculations [Fig 6A, (i) PRDM9]. Aggregation plots across PRDM9 binding sites showed that TH2BS12P is strongly enriched at the vicinity of PRDM9 containing sites. We found co-occupancy of TH2BS12P and PRDM9 to be highest at proximal distances suggesting that this histone mark is linked to genomic regions containing PRDM9. As the relative distance of TH2BS12P increases from PRDM9 binding site, there is decrease in overlap between TH2BS12P and PRDM9. Consistent with earlier observations, we found that majority of TH2BS12P reads were concentrated at H3K4me3 containing regions in relation to relative distance calculation as shown in Fig 6A, (ii) H3K4me3. The same was true for increased localisation of TH2BS12P at H3K4me3 marks associated with gene promoters and enhancer marks [Fig 6A, (iii) common H3K4me3].As the relative distance increases, the overlap of both these two histone marks also decreases. Furthermore, with respect to ‘B6 specific’ H3K4me3 marks, representing the recombination hotspots, we found increased overlap with the respect to distance of total TH2BS12P peaks [Fig 6A, (iv) B6 specific H3K4me3]. This demonstrates that the majority of occupancy of TH2BS12P modification is towards regulatory regions of the genome like gene promoters, enhancers etc that are positive for H3K4me3 marks. It is noteworthy from aggregation plot datasets that the fraction of TH2BS12P associated with total H3K4me3 was higher as compared to that of PRDM9. These results is highly consistent with the peak comparison data wherein we found ~48% association with total H3K4me3 containing regions as opposed to localisation to ~13% of recombination hotspots.

**Fig 6.**
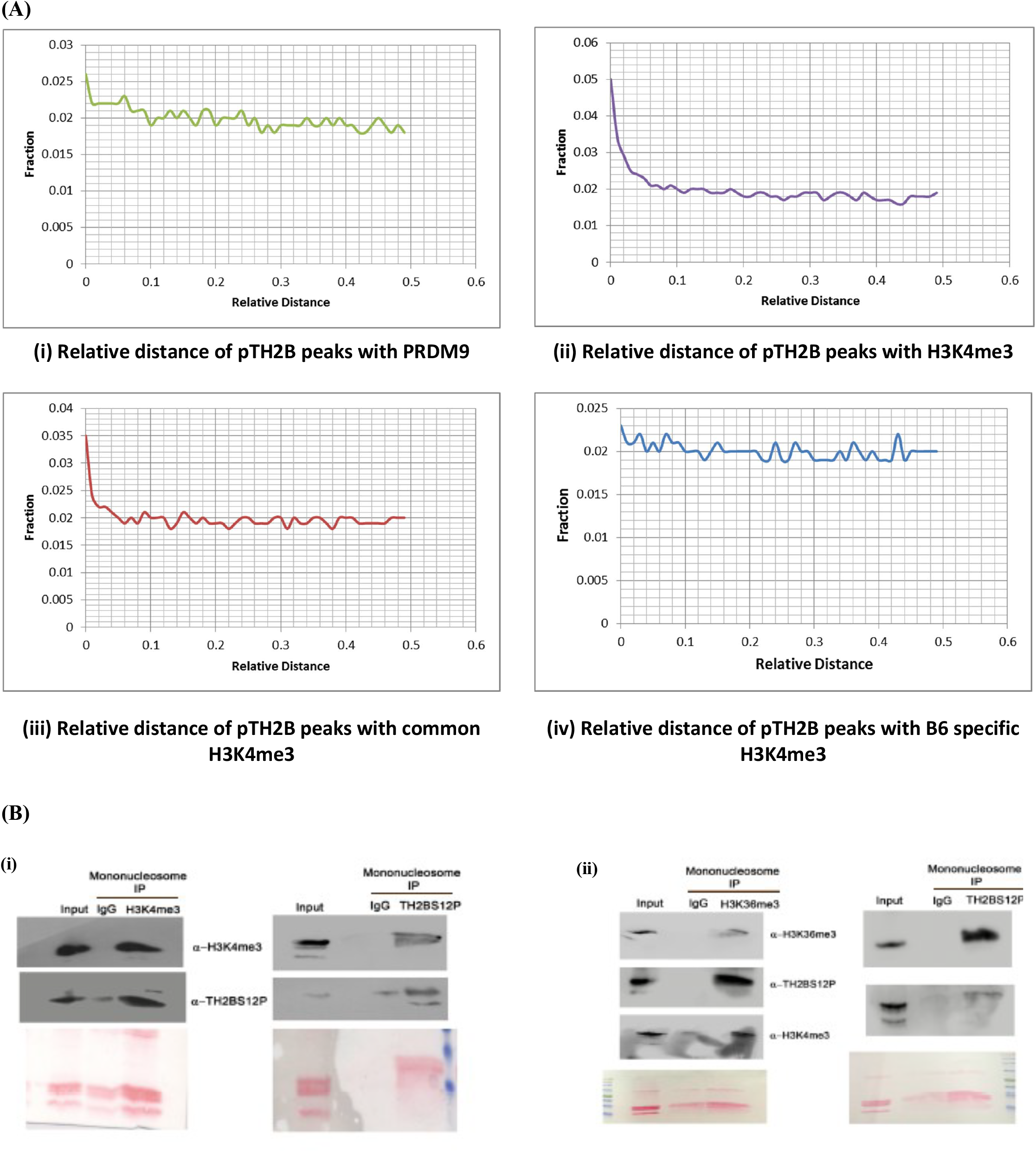
TH2BS12P histone mark is predominantly associated with genomic regions containing the histone mark H3K4me3-. **(A)** Aggregation plots showing the fraction of TH2BS12P reads with regard to relative distance of **(i)** PRDM9, **(ii)** total H3K4me3, (Hotspots), **(iii)** common H3K4me3 (present at TSS, enhancers) and **(iv)** B6 Specific H3K4me3; **(B)** Combinatorial histone marks in TH2BS12P containing mononucleosomes-Mononucleosome IP studies to determine the coexistence of transcription & recombination associated **(i)** H3K4me3 and **(ii)** H3K36me3 marks with TH2BS12P histone modification. Ponceau stained blots have been given for reference. The first lane in all the blots represents the Input, second lane-immunoprecipitation with the Rabbit IgG antibody, third lane refers to the IP using the specific antibody. The alpha alongside the blot refers to the antibodies that were used for western blotting.

Open chromatin regions like transcription start sites are initiation sites for meiotic recombination in yeast. However, hotspots and TSS are distinctly located in the mouse genome, also have similar histone mark profile like H3K4me3, H3K36me3 etc^62,64,71,73^. Peak overlap studies and aggregation plots had indicated the strong association of TH2BS12P with important histone marks like H3K4me3 and H3K36me3. To verify these results further, we therefore carried out immunoprecipitation reactions with mononucleosomes to determine the coexistence of transcription and hotspot associated histone marks H3K4me3 and H3K36me3 in a single nucleosome core particle. We found that H3K4me3 is associated with TH2BS12P containing mononucleosomes (Fig 6B (i), TH2BS12P IP). Conversely, co-IP with H3K4me3 antibody pulled down TH2BS12P protein (Fig 6B (i), H3K4me3 IP). To establish the association of TH2BS12P mark with H3K36me3, we also carried out immunoprecipitation studies with H3K36me3 antibody, wherein where we observed that TH2BS12P histone mark is associated with H3K36me3 containing mononucleosomes (Fig 6B (ii), H3K36me3 IP). Further, we also demonstrate the interaction of H3K36me3 and TH2BS12P IP by reciprocal IP (Fig 6B (ii), TH2BS12P IP). The combination of forward and reciprocal immunoprecipitation reactions with mononucleosomes proved further the association of TH2BS12P with both H3K4me3 and H3K36me3 marks *in vivo*.

For examining the possible presence of TH2BS12P modification at important genomic regions and for confirmation of the ChIP-sequencing dataset, we designed primers for various chromosomal loci across the mouse genome. The genomic regions with the input and IP tracks that have been selected for experimental validation have been shown in Fig 7A. By ChIP-PCR in P12 mouse testicular cells, we show the occupancy of TH2BS12P by at all the selected genomic regions, further validating the ChIP-seq results (Fig 7B, loci 1-12). There was no enrichment of TH2BS12P in a selected region of chromosome 1 that was used as negative control (Fig 7B, locus 13).

**Fig 7.**
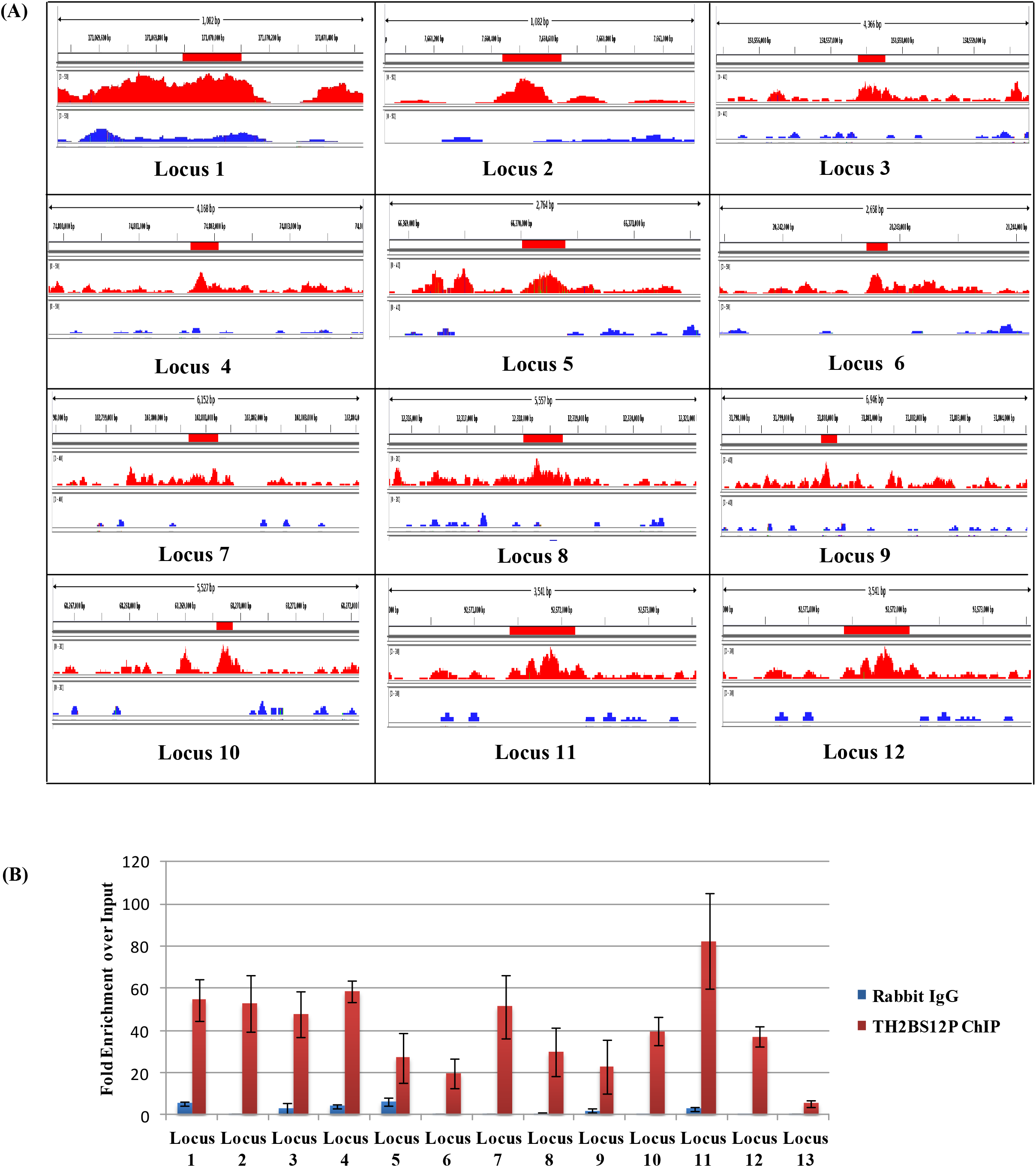
Validation of TH2BS12P ChIP-seq data by ChIP-PCR technique. **(A)** Genomic regions used for designing primers required to confirm the ChIP-sequencing dataset by ChIP-PCR studies. Visualization of selected TH2BS12P ChIP peaks showing the enrichment of TH2BS12P histone mark at the selected regions like locus 1(chr1:171067401-171072999), locus 2 (chr1:7658400-7662599), locus 3(chr2:158555400-158559799), locus 4(chr1:74799800-74803999), locus 5(chr14:66367200-66372399), locus 6(chr12:20240200-20245399), locus 7(chr8:102797400-102803599), locus 8(chr17:12315600-12321199), locus 9(chr15:31797600-31804599), locus 10(chr14:68268800-68269999), locus 11(chr3:92571400-92572199), locus 12(chrX:8074600-8074799), locus 13(chr1:132759100-132759222) using Integrative Genome Viewer (IGV). The y-axis represents the reads per million (rpm) values; The x-axis represents the selected genomic coordinates in base pairs; A specific region in chromosome 1(locus 13) was used as negative control based on absence of TH2BS12P peaks. The peaks in the input (marked in blue color) and IP track (marked in red color) are shown in the same scale for comparison. **(B)** Validation of TH2BS12P genome-wide occupancy data using ChIP-PCR technique which shows the enrichment of TH2BS12P histone mark at the selected loci across the mouse genome. A specific region in chromosome 1 was used as negative control for this study. The data are plotted in terms of fold enrichment values of TH2BS12P and Rabbit IgG against input control. ChIP-PCR experiments were done for three biological replicates including technical duplicates for a single biological replicate. The data are plotted in terms of mean ± S.D.

### TH2BS12P modification is associated with important proteins and histone marks related to transcription, meiotic recombination and pericentric heterochromatin formation

Since TH2BS12P is a nucleosomal histone protein, we were interested to determine the interaction protein partners that are physically associated with TH2BS12P containing mononucleosomes. Micrococcal nuclease digestion was used to obtain the mononucleosomes. The profile of fragments obtained with respect to time of MNase digestion has been shown in Fig S4C. To address this, we performed mass spectrometry after performing immunoprecipitation using the TH2BS12P antibody to identify the interacting proteins. From Fig 8A, we can see that TH2BS12P antibody is able to efficiently pull down proteins in comparison to non specific IgG control. We have further validated the specificity of the TH2BS12P antibody, where we show that the serine 12 phospho-peptide competes with the antibody in the immunoprecipitation reaction (Fig S1E). The interacting proteins were identified on the basis of enrichment of immunoprecipitated proteins with respect to their presence in non specific pre-immune IgG lane. Mass spectrometric analyses revealed important proteins associated with the functions of recombination and transcription as shown in the table (Fig 8B) further revalidating the ChIP-seq results where we observed the predominant association of TH2BS12P with important genomic regions related to gene regulation and recombination processes. The associated functions of each of these proteins have also been explained in the Fig 8B.

**Fig 8.**
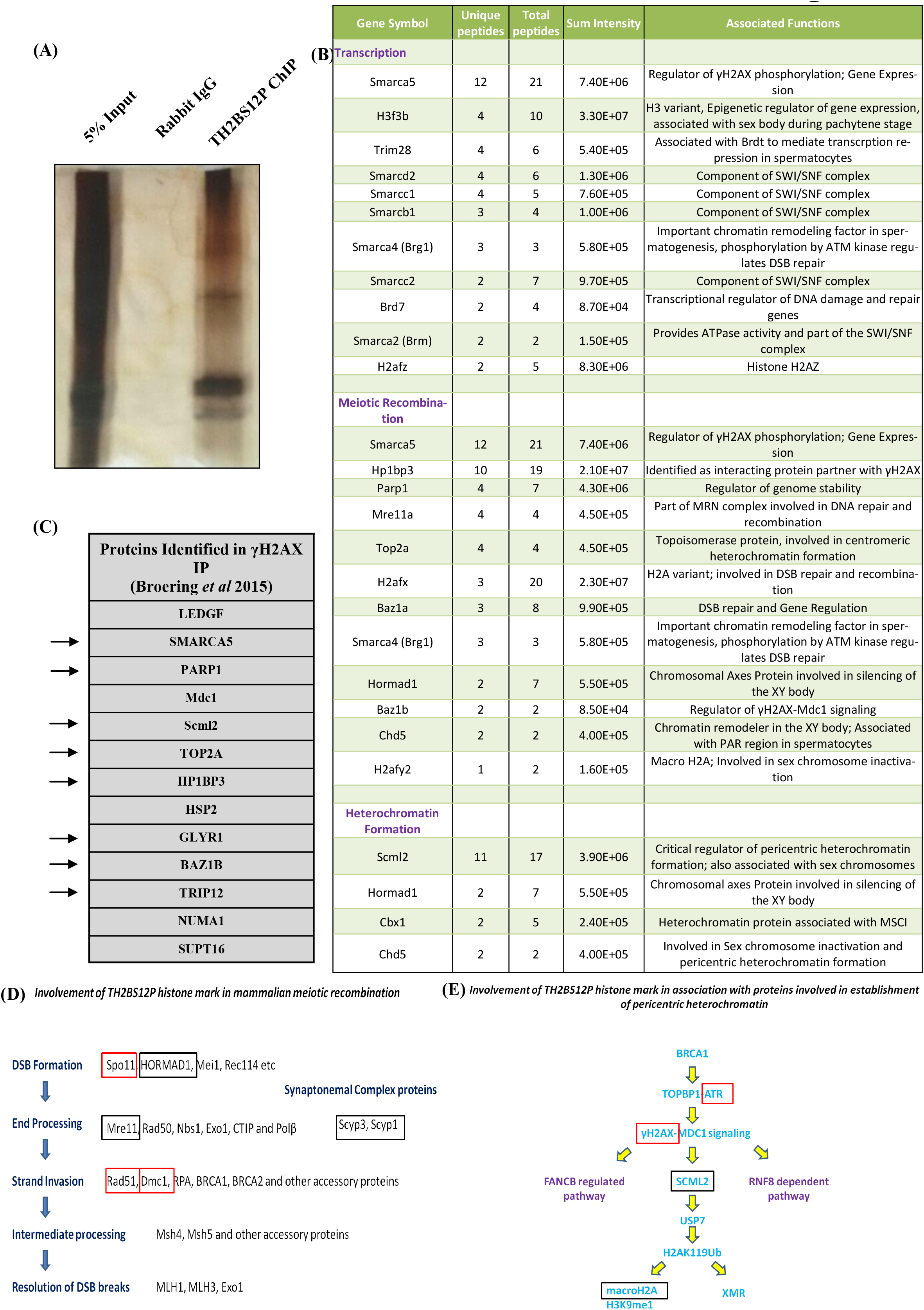
Identification of interacting protein partners and combinatorial histone marks of TH2BS12P containing nucleosomes-. **(A)** Silver Stained image of the TH2BS12P ChIP-Input lane refers to 5% input; Rabbit IgG lane refers to non specific IgG control; ChIP refers to TH2BS12P IP lane. **(B)** List of interacting proteins of TH2BS12P containing mononucleosomes as determined by mass spectrometry-The first row is the gene/protein name; The second and third rows refers to the number of unique and total peptides respectively (unique peptide sequences have been given in supplementary); the fourth row refers to the sum intensity that refers to the peak intensity values for all the peptides matched to a particular protein. The fifth row contains the known functions associated with those particular proteins. **(C)** List of proteins identified in γH2AX IP study as given the report published by Broering *et al* (2015). **(D)** Involvement of TH2BS12P histone marks in epigenetic pathways related to meiotic recombination and pericentric heterochromatin formation-The boxes in red highlight the proteins that were shown to be associated with TH2BS12B by immunofluorescence or immunoprecipitation assay whereas the boxes in black refers to proteins that interact with TH2BS12P as shown by their association by mass spectrometry technique.

Smarca5 is known to regulate Tyr142 phosphorylation on H2AX. It also associates with Cer2 to regulate gene expression^9^. Scp1, Scp2 and Scp3 are various components of the synaptonemal complex^1,39,43,65,85,88,89^. Hormad1 functions along with Spo11 and DSB machinery to regulate meiotic recombination^70^. Mre11 is part of the MRN complex that is involved in end resection^12^. Trim28 is known to associate with Brdt to mediate transcriptional repression in spermatogenic cells^83^. In ESCs, H3.3 has a rapid turnover at promoters and enhancers that possess various active histone marks like H2AZ, H3K27ac etc^26^. Interestingly, various important proteins involved in transcriptional regulation like H3.3, Brg1, H2AZ, were also identified as interacting proteins by mass spectrometry. Npm1 is a known histone chaperone. Consistent with the localization of TH2BS12P in the XY body, many proteins associated with the XY body functions like Scml2^20,36^, Cbx1^34^, macroH2A^58^, H2AX^14^ were also identified in the mass spectrometric analyses. Scml2 is a germline specific polycomb protein, associated with silencing of somatic or progenitor gene expression program in spermatocytes. It is also a critical regulator of epigenetic programming in sex chromosomes. Scml2 works downstream of γH2AX-Mdc1 signaling^20,36^(Fig 8E). The observation that we found TH2BS12P to be associated with γH2AX in mononucleosomes, gives a major indication of TH2BS12P function in regulating γH2AX-Mdc1-Scml2 axis in the XY body. Synaptonemal complex proteins are implicated in proper crossover formation and completion of meiosis. Also, synaptonemal complex proteins like Scp1, Scp2 and Scp3 were found to be associated with TH2BS12P mononucleosomes. Apart from this, chromosomal axes proteins like HORMAD1 that are involved in checkpoint surveillance required for meiotic progression is also associated with TH2BS12P histone mark. It is also known that Cbx1/HP1β and macroH2A are co-associated with the PAR region during male meiosis^79^. This reconfirms the association of TH2BS12P with the PAR region as both these proteins also shown to interact with TH2BS12P containing mononucleosomes (Fig 8B). In this list, we also obtained core nucleosomal histones, thus serving as internal control for the isolation of mononucleosomes.

In comparison to the proteins that were shown to be associated with γH2AX containing nucleosomes in a another report^9^, we find that many proteins (highlighted by arrows) were common and also associated with TH2BS12P containing mononucleosomes. This suggests that both these histone marks have similar interacting protein profile. These suggest that TH2BS12P and γH2AX co-associate to recruit similar set of proteins in turn to regulate DSB processes in spermatocytes. The important point to be kept in mind is also that TH2BS12P histone mark also interacts with proteins related to transcriptional regulation as shown by mass spectrometry. This indicates that TH2BS12P associates with different set of protein machineries in turn regulating different loci in a context-dependent manner. Taken together, a recurrent theme that emerges from our findings is that the novel histone mark TH2B Serine 12 phosphorylation is predominantly associated with H3K4me3 containing genomic regions and protein players involved in processes such as transcriptional regulation, recombination hotspots and pericentric heterochromatin in mammalian spermatocytes.

## Discussion

Histone tails of H2B do not exhibit specific structures in high resolution crystal structures. They contribute to the stabilisation of chromatin structures through contacts with DNA backbone and acidic patch of adjacent nucleosomes. Further, tail domains provide the stabilisation effect by suppressing the accessibility of the DNA and regulating nucleosome mobility^2,52^. In order to understand the unique functions of TH2B, it becomes important to study the function of the amino terminal tail of TH2B in terms of understanding the biological role of post-translational modifications and contribution of amino acid sequence differences to the overall nucleosome structure. Although there are some unique PTMs in H2B and TH2B in the globular domain and C-terminal tails^51^, we surmised that the unique N terminal sequence of TH2B and possible PTMs may contribute to biological functions of TH2B in testicular germ cells. We have recently described the complete repertoire of TH2B PTMs from pachytene spermatids and round spermatids^51^. Our results to the mass spectrometric analysis of TH2B PTMs using a modified procedure of enzyme digestion and processing of peptides revealed serine 12 phosphorylation in TH2B. This serine is highly conserved in most of the mammals suggesting possible role(s) in germ cells. Therefore, we focussed our studies on understanding the biological function of TH2BS12P modification in the context of processes characteristic of meiotic prophase I in rodent spermatocytes.

The immuno-staining of TH2BS12P in the pachytene spermatocyte revealed an interesting phenomenon in that this modification was highly enriched in the axes of the XY body apart from the staining that is observed outside the XY body. The intense staining in the XY body axes could be because of the increased DSB density that occurs in the XY body compared to autosomal DSBs. Importantly, a specialised chromatin configuration is found in the XY body where the DNA is organized into longer axes and shorter loops promoting higher DSB density compared to autosomes^23^. This could suggest that TH2BS12P may aid in formation of this specialised chromatin structure in the PAR region. Due to differences in chromatin configuration that is observed in PAR versus autosomal hotspots, TH2BS12P could play context-specific roles in autosomal and PAR DSB formation wherein other important players of recombination or histone marks could act in concert in determining the specific hotspot activation.

ATM and ATR signaling pathways are associated with autosomal and XY body DSBs respectively in spermatocytes. We were therefore first interested in understanding the association of TH2BS12P with recombination proteins like Spo11, pATM, ATR etc on the basis of dense localisation of this histone mark at the axes of the sex body of pachytene spermatocyte. We observe that Spo11 is associated with TH2BS12P in the leptotene/zygotene intervals where autosomal DSBs occur. Apart from the role of TH2BS12P in the ATM dependent signaling in autosomal DSBs, we show that TH2BS12P is involved in ATR mediated signaling in the sex body DSB formation. Furthermore, we show that the marker proteins of DSB initiation and strand invasion Spo11 and Rad51 associated chromatin fragments are associated with TH2BS12P histone mark. Since it is known that DSB machinery is recruited to the axes, wherein adaptor proteins are postulated to play a major role in promoting axis-loop interaction^17^, the localization of this modification in the context of binding sites in the genome had to be determined by ChIP-sequencing analyses. ChIP-sequencing analyses of genome wide distribution of TH2BS12P gave us a comprehensive picture of occupancy of this modification in relation to binding sites of important recombination and transcription related proteins. We found only a subset of recombination hotspots (~13.48%) were associated with TH2BS12P. It is pertinent to note that recombination hotspots and gene promoters share similar histone mark profile like H3K4me3 and H3K36me3 but the formation of which are catalysed by different enzymes. Apart from localisation at H3K4me3-containing hotspots, we prove that the majority of TH2BS12P interacting regions was linked to other H3K4me3 containing regions (48.5%) like gene promoters, enhancers etc. Aggregation plots also suggested that the majority of TH2BS12P reads were linked to H3K4me3 histone marks excluding the hotspots. It is important to note that ChIP sequencing were carried out in P12 testicular cells, therefore, the association of TH2BS12P with XY body might be underrepresented in P12 mouse testicular cells. For this, a specific ChIP-seq in P21 cells would give a clearer picture of involvement in XY body DSBs and repair phenomena.

HBR domain in core histone H2B has been shown to regulate nucleosome mobility and DNA repair^59^. TH2BS12P could modulate HBR domain in terms of structural implications or also could crosstalk with specialised HBR domain PTMs to regulate chromatin structure and DNA repair. Also, chromatin relaxation in response to DSB breaks is mediated by ATM signaling pathway in somatic cells^84^. This could mean that the TH2B mediated nucleosomal instability in combination with the phosphorylation moiety could contribute to this phenomenon in subset of selective recombination hotspots. The modification resides in the disordered amino terminal tail of TH2B, which means that it plays an important role more in mediating protein-protein interactions than protein-DNA interactions. More importantly, immunoprecipitation assays showed the association of TH2BS12P with recombination related histone marks like H3K4me3, H3K36me3 and DSB initiator γH2AX in mononucleosomes (shown in Figure 7D). ‘Histone code hypothesis’ proposes the role of histone modifications acting in concert or sequentially to bring about chromatin mediated DNA-transaction processes like transcription, DNA repair etc^22^. This could mean that TH2BS12P along with other histone modifications like H3K4me3, H3K36me3, γH2AX within those particular nucleosomes could either contribute to higher order chromatin structure necessary for hotspot activation or a signature for recruiting the appropriate protein machinery on the chromatin template. Based on this, there are three possible interdependent functions of TH2BS12P histone mark-(a) TH2BS12P could regulate HBR domain structure and regulate crosstalk with HBR domain PTMs; (b) TH2BS12P could associate with γH2AX and PRDM9-mediated-H3K4me3 marks to regulate recombination events in spermatocytes; (c) TH2BS12P could function in H3K4me3 mediated recruitment of effector proteins and chromatin modellers to determine transcription functions. Thus, TH2BS12P histone mark could play a major role in creation of a chromatin environment along with other histone marks to create platform for transcription, enhancer function or Spo11-mediated DNA break formation.

Mass spectrometry analysis of interacting proteins of TH2BS12P containing mononucleosomes also revealed the signaling pathways of the involvement of this modification in context of transcription, meiotic recombination and pericentric heterochromatin formation. In regard to transcription, we found interaction of TH2BS12P with transcription proteins like H2AZ, Baz1a, Brg1, SWI-SNF components etc. In context of recombination proteins, we found association with HORMAD1 that is linked to DSB initiation and Mre11 which is associated with DNA resection processes (Fig 8D). It is to be noted that the immunoprecipitation assays for mass spectrometry studies were performed with mononucleosomes, it is therefore possible that the tightly bound proteins to mononucleosomes would only be found in mass spectrometry results. We also observed interaction of TH2BS12P with Scml2 by mass spectrometry, which is the critical regulator of pericentric heterochromatin in mammalian spermatocytes. ATR dependent signaling operates in the recombination processes in the XY body, leads to γH2AX-mediated Mdc1 signaling. The downstream effector, Scml2 is the major protein involved in heterochromatin formation wherein it prevents RNF2 mediated ubiquitination of H2A on sex chromosomes^20,36^. This pathway leads to macroH2A deposition via recruitment of USP7 enzyme. Also, it marks somatic genes on autosomal for repression. Apart from the SCML2 dependent pathway, we did not find association with FANCB and RNF8 pathway associated proteins (Fig 8E). Mass spectrometric analyses therefore revealed the association of TH2BS12P containing mononucleosomes towards important protein players involved in pericentric heterochromatin formation, transcription and meiotic recombination processes (8D and 8E). This study also raises an interesting question about what histone or histone marks and their PTMs combinations define a ‘TSS’ or ‘hotspot’ in meiotic spermatocytes.

In a previous study, Montellier *et al*^41^ used two mouse models to explain the function of TH2B in spermatogenesis. The loss of TH2B (knockout mouse model) was compensated by upregulation of H2B and additional PTMs were acquired on the core histones H2B, H3 and H4. Specialised modifications like lysine crotonylation were observed on H2B to complement the functions of TH2B. From the tagged-TH2B model, it was proven that the tag interfered with the function of TH2B creating a dominant negative phenotype. The tagged TH2B expressing mice were sterile with defects in histone to protamine replacement. TH2B, thus was proven to play a major role in processes related to histone to protamine replacement in spermiogenesis. This report reveals for the first time the association of a TH2B modification in DSB repair and recombination processes characteristic of meiotic prophase I. In addition, the influence of TH2BS12P on 3D chromatin architecture influencing chromatin-templated events cannot be discounted.

## Materials and Methods

### Materials

All fine chemicals were obtained from Sigma Chemicals (USA) unless mentioned otherwise. Synthetic peptides were outsourced from Stork Bio Laboratories (Estonia). The secondary antibodies Donkey anti-mouse, Goat anti-rabbit, Goat anti-rabbit conjugated to Alexa dyes were obtained from Invitrogen (USA). Male Wistar rats and C57BL6 mice were obtained from the Animal Facility, JNCASR. All procedures for handling animals have been approved by the animal ethics committee of the centre.

### Purification of *in vivo* TH2B protein

The histones were isolated from 30-35 day old rat testes by acid extraction method. *In vivo* TH2B were purified by RP-HPLC technique using the published protocol^51^.

### Mass spectrometry for identification of posttranslational modifications on TH2B

#### (i) In-Gel Digestion

Gel bands were cut into one mm^3^ piece and washed twice with MilliQ water. The gel was destained using 1:1 methanol:50 mM ammonium bicarbonate for 1 min, twice. The gel pieces were dehydrated for 5 min using 1:1 acetonitrile: 50 mM ammonium bicarbonate followed by acetonitrile for 30 s. The gel pieces were dried in a speed-vac (Thermo Savant) for 10 min. The gel pieces were rehydrated in 25 mM dithiothreitol, 50 mM ammonium bicarbonate and incubated at 56 C for 20 min. After discarding the supernatant, the gel pieces were incubated in 55 mM iodoacetamide at RT for 20 min in the dark and subsequently were washed twice with water, dehydrated and dried as before. The dried gel pieces were rehydrated in 50 mM ammonium bicarbonate containing 250 ng of mass spectrometry grade trypsin (Promega) and incubated overnight at 37 C. Following digestion, the reaction mixture was acidified with 1% acetic acid and dried in a speed-vac to reduce the volume to 5 μL, to which 10 μL of mobile phase A was added for direct loading for LC-MS/MS analysis.

#### (ii) Liquid Chromatography-Tandem Mass Spectrometry

Each reaction mixture was analyzed by LC-MS/MS. LC was performed on a Easy nanoLC II HPLC system (Thermo). Mobile phase A was 94.5% MilliQ water, 5% acetonitrile, 0.5% acetic acid. Mobile phase B was 80% acetonitrile, 19.5% MilliQ water, 0.5% acetic acid. The 120 min LC gradient ran from 2% B to 35% B over 90 min, with the remaining time used for sample loading and column regeneration. Samples were loaded to a 2 cm x 100 um I.D. trap column positioned on an actuated valve (Rheodyne). The column was 13 cm x 100 um I.D. fused silica with a pulled tip emitter. Both trap and analytical columns were packed with 3.5 μm C18 resin (Zorbax SB, Agilent). The LC was interfaced to a dual pressure linear ion trap mass spectrometer (LTQ Velos, Thermo Fisher) via nano-electrospray ionization. An electrospray voltage of 1.8 kV was applied to a pre-column tee. The mass spectrometer was programmed to acquire, by data-dependent acquisition, tandem mass spectra from the top 15 ions in the full scan from 400 - 1400 m/z. Dynamic exclusion was set to 30 s.

#### (iii) Data Processing and Library Searching

Mass spectrometer RAW data files were converted to MGF format using msconvert. Detailed search parameters are printed in the search output XML files. Briefly, all searches required strict tryptic cleavage, 0 or 1 missed cleavages, fixed modification of cysteine alkylation, variable modification of methionine oxidation and expectation value scores of 0.01 or lower. MGF files were searched using X! Hunter against the latest library available on the GPM at the time. Other searches used the cRAP contaminant library from the GPM and libraries constructed from the most recent ENSEMBL release available at the time. MGF files were searched using X!! Tandem using both the native and k-score5 scoring algorithms and by OMSSA. All searches were performed on Amazon Web Services-based cluster compute instances using the Proteome Cluster interface. XML output files were parsed, and non-redundant protein sets determined using in-house scripts. Proteins were required to have 2 or unique peptides with E-value scores of 0.01 or less, 0.001 for X!Hunter and protein E-value scores of 0.0001 or less.

### Alignment of the amino acid sequences

The amino acid sequences were obtained for TH2B from Uniprot and p-blast was performed to identify plausible protein counterparts in other mammalian species. Multiple sequence alignment (MSA) was performed for selected mammals for TH2B to elucidate the sequence conservation across species.

### Antibody Generation

Peptides corresponding to TH2BS12P modification (CKGTTI(pS)KKGFK), H2B(KSRPAPKKGSK) were injected into rabbits, and the 14-day cycle of antibody generation was followed. Immunoglobulins were purified by caprylic acid based purification. Peptide-affinity based purification with the Sulfolink columns containing immobilized peptides, was used to purify the TH2BS12P and H2B specific antibodies. The TH2BS12P antibody was outsourced from Abgenex company.

### Preparation of nuclear lysates

Nuclear lysates and chromatin fraction were prepared by the method described previously with modifications^72^. Briefly, testes were dissected in cytoplasmic lysis buffer (10mM HEPES pH 7.5, 50mM NaCl, 0.5M sucrose, 0.5% Triton-X-100, 0.1mM EDTA, 1mM DTT, protease inhibitor cocktail), incubated on ice for 15 minutes and centrifuged at 1500g for 7 minutes. The nuclear pellet was resuspended in Buffer B1 (10mM HEPES pH 7.5, 500mM NaCl, 0.1mM EDTA, 1mM DTT, 0.5% NP-40, protease inhibitor cocktail) to obtain nuclear lysates or Buffer B2 (10mM HEPES, 200mM NaCl, 1mM EDTA, 0.5% NP-40, protease inhibitor cocktail) for isolation of chromatin. The nuclear lysates were clarified by centrifugation at 15100 X g for 10 minutes. After incubation on ice for 15 mins in Buffer B2, the pellet was recovered by centrifugation at 17760 X g for 2 minutes. Further, the pellet was suspended in Buffer C (10mM HEPES, 500mM NaCl, 1mM EDTA, 1% NP-40, protease inhibitor cocktail), sonicated for 15 cycles and chromatin fraction was obtained after centrifugation at 17760 X g for 1 minute.

### ELISA

Peptides were used at 200 ng per well. The pre-bleed and immune sera were used at 1:5000 dilution. Goat anti-rabbit HRP were used as the secondary antibody at 1:5000 dilution. TMB (3, 3’, 5, 5’-Tetramethylbenzidine) was used as the substrate for the reaction. After three minutes of enzyme-substrate reaction, the plate was read at 450 nm.

### Dot Blot Analysis

Two μg of peptides corresponding to TH2B (CKGTTISKKGFK), TH2BS12P (CKGTTI(pS)KKGFK),H2B(KSRPAPKKGSK),H2BS14P(SRPAPKKG(pS)K KC) were applied as separate spots on the nitrocellulose membrane. After drying, the blot was subjected to steps of western blotting with the TH2BS12P antibody.

### Preparation of meiotic spreads

Meiotic spreads were prepared using the published protocol^81^. Briefly, testes were decapsulated and chopped in PBS solution (pH 7.4). The cell pellet was resuspended in hypotonic buffer (30mM Tris, 17mM sodium citrate, 50mM sucrose, 5mM EDTA, 0.5mM DTT, protease inhibitor cocktail) and incubated for 30 mins. The pellet was resuspended in 100 mM sucrose solution, and the nuclei were spread onto PFA-coated slides. The slides were kept for drying at room temperature for 2 hours and proceeded for immunofluorescence studies.

### Immunofluorescence

The slides were kept in blocking solution (3% BSA solution in PBS) for 1 hour at room temperature, then treated with primary antibody overnight in the cold room, washed with 0.1% PBST solution and then incubated with secondary antibody for 1 hr at room temperature. Next, washes were given with PBST solution and the smears were mounted using DAPI solution. Images were acquired by Zeiss confocal laser scanning microscope (LSM880 or LSM510). Zen software was used for image analysis. For the analysis, each image was subjected to background correction and threshold calculation. After setting the threshold, colocalization percentages were calculated taking to account pixel counts in the three quadrants given in the formula below. All data were confirmed with at least three or four independent mice and rats.

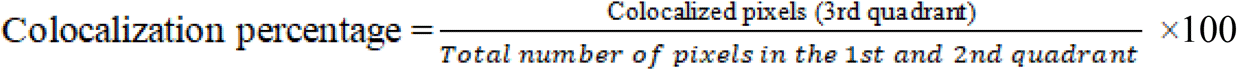

### Isolation of mononucleosomes

Immunoprecipitation using mononucleosomes were carried out using the published protocol^41^. Briefly, the rat testes were dissected and homogenised in lysis buffer (60mM KCl, 15mM NaCl, 15mM Tris-HCl, 0.03% Triton-X-100, 0.34M Sucrose, 2mM EDTA, 0.5mM EGTA, 0.65mM spermidine, 1mM DTT, 1% glycerol, protease and phosphatase inhibitor cocktail), centrifuged at 650 X g for 10 mins at 4°C. The pellet was washed with wash buffer containing 60mM KCl, 15mM NaCl, 15mM Tris-HCl, 0.34M Sucrose, 0.5mM EGTA, 1mM DTT, 0.5mM PMSF and protease and phosphatase inhibitor cocktail. The pellet was resuspended in MNase buffer (10mM Tris-HCl, 10mM KCl, and 2mM CaCl_2_). The nucleosome fraction was isolated by centrifugation at 650 X g for 10 mins at 4°C, mixed with LSDB250 buffer (20% glycerol, 50mM HEPES, 3mM MgCl_2_, 250mM KCl, protease and phosphatase inhibitor cocktail) and proceeded with the immunoprecipitation protocol.

### ChIP sequencing of TH2BS12P modification in P12 mice

DNA was isolated from TH2BS12P immunoprecipitated mononucleosomes by phenol-chloroform method. The quality control of the DNA samples was done using the Qubit and Tapestation methods. The libraries were subjected for 40 million depth paired-end (100 bp x 2) sequencing that was carried using Illumina HiSeq 2500. FASTQ files were obtained and data analyses were carried out further.

### Data Analysis

FASTQ files were aligned to mm10 genome assembly using Bowtie2^31^. While aligning, unpaired and discordant reads were removed. The aligned files were sorted and indexed accordingly, and also made free from PCR duplicates. The sorted aligned files of Input and antibody treated (IP) were merged respectively using Samtools Merge^53^. The peaks were called between the merged input and IP files using SICER 1.1 Version^67^ with the following parameters-Redundancy threshold: 1; Window size: 200bp; Fragment size: 150bp; Gap size: 600bp; FDR: 0.01. The final peaks were shortlisted giving the cutoff of >1.5 fold change. The overlapping peaks with the respective protein and histone marks were identified using Bedtools Intersect. Final peaks were annotated using HOMER and repeats were classified using the repeat data from UCSC Table Browser.

### Aggregation Plot

The relative distance between TH2BS12P peaks and the earlier mentioned protein and histone marks were computed using reldist of Bedtools suite considering +/-2Kb from the peak summits and were plotted in R.

### Immunoprecipitation and Quantitative PCR

The chromatin or mononucleosomes fraction was incubated with the given antibody for overnight at 4°C. Protein A or Protein G dynabeads were added the next day, washed with Buffer C (with 150mM NaCl) for ChIP. LSDB250 buffer was used as the wash buffer for immunoprecipitation studies with mononucleosomes. After the washes, the beads were either proceeded with DNA extraction for PCR analysis or boiled in 5X SDS dye for western blotting. After washing of beads, DNA was eluted from the beads as follows-210 μl of the elution buffer was added, incubated at 65°C overnight for de-crosslinking. 200 μl of TE buffer was added the next day, subjected for RNase (Final; 40μg/ml) and proteinase K (Final; 100μg/ml) treatment and DNA were extracted by phenol-chloroform method. DNA that was purified from TH2BS12P ChIP were proceeded for ChIP-seq analyses. SYBR kit from TAKARA was used to set up quantitative PCR reactions. PCR was carried out for 40 cycles and was followed by melt curve analyses before recording the raw Ct values. The fold enrichment values were calculated over input taking the percentage of input used for the ChIP procedure and the Ct values obtained for the target genomic region from Input and ChIP DNA. PCR was carried out in duplicates for each of the three biological replicates.

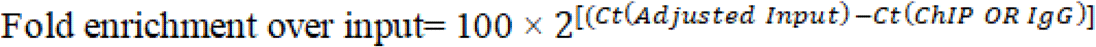

### Primer design for ChIP-PCR studies

Peak summits corresponding to high TH2BS12P occupancy like intergenic, intronic, hotspots, promoters etc. were chosen for experimental validation using ChIP-PCR. In order to maintain the rigour of primer design, primers were designed using the Primer BLAST and primer 3 tools and were also verified computationally using NCBI Primer Blast and UCSC In-silico PCR. Verification of primer dimer formation were also considered during the design.

### Mass spectrometric identification of interacting protein partners of TH2BS12P containing mononucleosomes

Immunoprecipitation of TH2BS12P containing proteins were carried out and the proteins were extracted from the beads using the elution buffer of the Pierce co-IP kit. The eluted proteins were resolved on 15% SDS gel and the gel was subjected to Coomassie staining. The stained wells corresponding to the IgG and IP lanes were sent further for mass spectrometry to determine the interacting proteins.

#### (i) Methods for Protein Sequence Analysis by LC-MS/MS

Excised gel bands were cut into approximately 1 mm^3^ pieces. Gel pieces were then subjected to a modified in-gel trypsin digestion procedure^66^. Gel pieces were washed and dehydrated with acetonitrile for 10 min. followed by removal of acetonitrile. Pieces were then completely dried in a speed-vac. Rehydration of the gel pieces was with 50 mM ammonium bicarbonate solution containing 12.5 ng/μl modified sequencing-grade trypsin (Promega, Madison, WI) at 4°C. After 45 min., the excess trypsin solution was removed and replaced with 50 mM ammonium bicarbonate solution to just cover the gel pieces. Samples were then placed in a 37°C room overnight. Peptides were later extracted by removing the ammonium bicarbonate solution, followed by one wash with a solution containing 50% acetonitrile and 1% formic acid. The extracts were then dried in a speed-vac (~1 hr). The samples were then stored at 4°C until analysis.

On the day of analysis the samples were reconstituted in 5 - 10 μl of HPLC solvent A (2.5% acetonitrile, 0.1% formic acid). A nano-scale reverse-phase HPLC capillary column was created by packing 2.6 μm C18 spherical silica beads into a fused silica capillary (100 μm inner diameter x ~30 cm length) with a flame-drawn tip^50^. After equilibrating the column each sample was loaded via a Famos auto sampler (LC Packings, San Francisco CA) onto the column. A gradient was formed and peptides were eluted with increasing concentrations of solvent B (97.5% acetonitrile, 0.1% formic acid).

As peptides eluted they were subjected to electrospray ionization and then entered into an LTQ Orbitrap Velos Pro ion-trap mass spectrometer (Thermo Fisher Scientific, Waltham, MA). Peptides were detected, isolated, and fragmented to produce a tandem mass spectrum of specific fragment ions for each peptide. Peptide sequences (and hence protein identity) were determined by matching protein databases with the acquired fragmentation pattern by the software program, Sequest (Thermo Fisher Scientific, Waltham, MA)^13^. All databases include a reversed version of all the sequences and the data was filtered to between a one and two percent peptide false discovery rate along with filter to being set to at least 1 unique peptide per protein.

### Western Blot Analysis

For western blot, the proteins were resolved by SDS-PAGE gel electrophoresis and then transferred onto a nitrocellulose membrane using the semi-dry transfer technique. The membrane was blocked using 5 % skimmed milk or 3 % BSA (diluted in TBS) for 1 hr at room temperature, then incubated with the specific primary antibody for 1 hr at room temperature or overnight at 4°C. The blots were given multiple washes with 0.1% PBST or TBST for 10 min each. Next, the blot was incubated with the secondary antibody (anti-rabbit /anti-mouse) for 1 hr at room temperature. Membranes were washed extensively with 0.1% PBST or TBST and developed using the ECL kit (Thermo Scientific). For the peptide competition assay, fifty fold molar excess of the modified and unmodified peptides were added to the antibody solution and mixed for 3 hrs at 4°C before the addition to the blot.

## DATA Access

The ChIP-sequencing dataset containing the raw and processed BED files have been deposited in the Gene Expression Omnibus (ID-Once Available, process has been initiated). The tables of primer sequences and antibodies used have been given in the Supplemental Materials.

List of DATASETS used for computational Data Analyses

**List of DATASETS used for computational Data Analyses**
| **Dataset** | **Geo Accession IDs** |
| --- | --- |
| DSB Hotspots | GSE93955 |
| PRDM9 | GSE93955 |
| H3K4me3 | GSE35498 |
| H3K4me3 common | GSE93955 |
| H3K4me3 (B6 specific) | GSE93955 |
| H3K36me3 | GSE93955 |
| Dmc1 | GSE93955 |
| H4K8ac | GSE77277 |

## Acknowledgements

We thank Dr. Brian Balgely of Bioproximity for mass spectrometry. We acknowledge Dr. R.G Prakash of the Animal Facility and Suma B.S with Sunil Kumar of the Confocal Imaging Facility at JNCASR. We also thank members of the Imgenex facility for carrying out the antibody generation project. We also thank Tomaino Ross of TAPLIN Mass spectrometry facility at Harvard for mass spectrometry based identification of interacting proteins. M.R.S. Rao thanks Department of Science and Technology, Government of India for J.C. Bose and SERB Distinguished Fellowships and this work was financially supported by Department of Biotechnology, Govt. of India (Grant Number BT/01/COE/07/09).

## Author Contributions

I.A.M, S.P, R.R performed the experiments. I.A.M, S.P and M.R.S designed the experiments and wrote the manuscript. U.B performed the computational data analysis associated with the ChIP seq dataset. All authors discussed the results and approved the final manuscript.

